# Differential Remodeling of the BRCA1 Interactome by a Unique Minimal Clinical Truncation Mutation

**DOI:** 10.1101/2024.11.26.625539

**Authors:** Jingjing Chen, Mihaela E. Sardiu, Michaella J. Rekowski, Jules Nde, Zachary S. Clark, Shrikant Anant, Michael P. Washburn, Roy A. Jensen

## Abstract

BRCA1, a critical tumor suppressor gene, plays an essential role in maintaining genomic stability through its involvement in DNA double-strand break repair, particularly via homologous recombination. Loss or impairment of BRCA1 function disrupts this repair pathway, resulting in genomic instability and significantly increased susceptibility to breast and ovarian cancers. To elucidate the molecular mechanisms by which BRCA1 mutations contribute to tumorigenesis, we employed quantitative mass spectrometry-based proximity labeling and affinity purification to identify cancer-specific protein-protein interactions (PPIs). Our integrated omics and visualization analysis of interactors revealed that the BRCA1-Y1853ter mutant, through its interaction with BARD1, perturbs the interactome and impacts cellular processes within the cytoplasm and nucleoplasm. Structural data further corroborated these findings, showing enhanced binding between the mutant BRCA1 and specific interactors, suggesting an altered functional profile. Together, these observations raise the hypothesis that the BRCA1-Y1853ter mutant may exhibit gain-of-function characteristics, providing new insights into the molecular and cellular effects of mutations in the BRCA1 C-Terminal (BRCT) domain and their implications for the pathogenesis of breast and ovarian cancers.

## 1. Introduction

BRCA1 functions as a pivotal tumor suppressor gene in the etiology of ovarian and breast cancers ^1–3^. Individuals harboring BRCA1 germline mutations exhibit a markedly increased susceptibility to ovarian cancer, with risks ranging from 39% to 46%, and a notably elevated predisposition to breast cancer, with probabilities ranging from 57% to 65% ^4–7^. Significantly, up to 80% of these cases manifest as the triple-negative subtype, characterized by the absence of estrogen receptor (ER), progesterone receptor (PR), and human epidermal growth factor receptor 2 (HER2) expression ^8,9^. This subtype of breast cancer is presently acknowledged as the most aggressive and refractory to conventional targeted therapies. Therefore, an in-depth investigation into the mechanistic basis of BRCA1 mutations driving triple-negative breast cancer (TNBC) is imperative ^10–12^.

BRCA1 encodes a large polypeptide consisting of 1863 amino acids. Its principal functional domains include the RING domain at the N-terminus, possessing E3 ubiquitin ligase activity involved in regulating ubiquitination and modification of histone H2A, claspin, RNAPII and other related substrates crucial in chromatin remodeling ^13^. Another domain is the C-terminal BRCA1 C-Terminal (BRCT) domain, that participates in homologous recombination-based DNA damage repair, G1/S and G2/M cell cycle regulation, transcriptional regulation, and signal transduction, essential for maintaining genome integrity and stability. Mutations in these domains can affect the biological functions of BRCA1, thus contributing to increased cancer susceptibility ^14^. The likelihood of cancer occurrence in BRCA1 carriers and the cancer subtype to some extent depends on the category and severity of BRCA1 mutations affecting its biological functions.

Residues aa 1642-1855 at the C-terminus of BRCA1 contain two tandem BRCT domains: BRCT-N (aa1642-1736) and BRCT-C (aa1756-1855). One hotspot for breast cancer clustering, c.5261 to c.5563 (aa1754-1855) (RHR=1.38; 95% CI, 1.22-1.55; P = 6 × 10^−_9^), falls within the BRCT-C domain ^15^. BRCA1 can specifically bind phosphopeptides containing the pSer-X-X-Phe motif through its BRCT-C domain, interacting with numerous phosphoproteins, including DNA endonuclease RBBP8/CtIP, carboxy-terminal helicase BACH1/BRIP1/FancJ, coiled-coil domain-containing protein 98 or CCDC98/ABRAXAS, forming three different complexes and executing its unique anti-tumor role in DNA repair and cell cycle modulation ^14,16^. BRCA1’s binding to DNA and non-phosphoproteins through the BRCT domain has also been reported ^17–19^.

Indeed, many representative clinical pathogenic BRCT mutants have been found to exhibit diminished or even abrogated binding with these key phosphoproteins, underscoring the pivotal role of these PPIs in anti-tumor mechanisms ^19,20^. Moreover, studies have indicated that pathogenic variants of BRCA1 in the BRCT domain display selective binding with specific interactors compared to the wild-type (WT) protein. Several BRCT variants have been documented to demonstrate significantly enhanced interaction with COBRA1/NELFB1 ^21^.

Recent findings have validated an augmented degree of binding between Hsp70, Hsp90, and BRCT variants, concomitant with the protein misfolding and functional deficiencies induced by mutations. Notably, 24 out of 31 pathogenic BRCA1-BRCT variants exhibited substantial interaction with HSPA8 (HSP70 homologues) and Hsp90AB1 (HSP90 homologues), whereas 9 out of 11 benign variants failed to interact with either chaperone ^22^. Computational modeling also suggests that mutations in BRCT could enhance the affinity of P53 binding and potentially disrupt the balance of P53-53BP1 interaction ^23^. Negative regulation of the AKT pathway was observed in BRCT mutant cells lacking the BRCT repeats ^24^. USP28 was also identified as having a higher binding affinity for BRCA1-5682insC ^25^. These research findings suggest that BRCT variants may manifest as potential gain-of-function mutations ^26^, potentially acquiring the capability to interact with common interactors via conformation change ^27,28^.

An additional crucial aspect to note is the indispensable role of BARD1 as an obligatory binding partner protein for BRCA1 in executing these vital anti-tumor functions, co-expressed across almost all tissues. The formation of a heterodimeric complex between BRCA1 and BARD1 occurs through their respective N-terminal RING domains, ensuring structural stability and functional interdependence. This complex not only augments the ubiquitin ligase activity of BRCA1 but also shields their individual nuclear export signals (NESs) by mutual binding, thereby preserving their nuclear functionality and promoting cell survival ^29^. Disruption of heterodimer formation leads to enhanced BRCA1 ubiquitination and subsequent proteasomal degradation, resulting in cytoplasmic sequestration of both proteins ^30–32^. Moreover, the BRCT domain of BARD1, which recognizes a distinct motif from the BRCA1 BRCT domain, facilitates targeted localization of the BRCA1/BARD1 heterodimer to DNA damage sites through its interaction with poly (ADP-ribose) (PAR) ^33^. These findings underscore the critical importance of investigating BRCA1 mutations within the context of heterodimer formation for elucidating carcinogenic mechanisms.

Liquid chromatography-tandem mass spectrometry (LC-MS/MS)-based proteomics provides an efficient and powerful tool for the systematic comparison and analysis of PPI networks across different conditions at the proteome-wide level^34^. In our study, employing a system ensuring the formation of heterodimers through co-expression of BARD1 to maintain stoichiometric balance, we utilized two state-of-the-art MS-based quantitative comparative interactome methodologies to scrutinize and contrast the interactomes of WT BRCA1 and BRCA1-Y1853ter variant- the smallest known penetrant germline BRCA1 truncation mutation ^35^. Its compact 11-amino acid deletion, involving the loss of only 3 highly conserved residues from the BRCT-C domain, highlights the minimal yet crucial nature of these key amino acids, making it an important model for investigating common pathogenic mechanisms among BRCT mutants. Both methodologies revealed that the BRCA1-Y1853ter mutation triggered a cascade of significant losses in PPIs, while also identifying mutation-specific interactions within the nucleus and cytoplasm. These findings offer insights into the molecular mechanisms underpinning BRCA1-Y1853ter mutation-driven carcinogenesis, potentially revealing mutation-specific PPIs indicative of a gain-of-function carcinogenic mechanism conferred by BRCA1 mutations. Moreover, the delineation of these unique PPIs may pave the way for the identification of potential therapeutic targets.

## 2. Materials and Methods

### 2.1 Cells and Cell Culture

The AD-X-293 cell line used for adenoviral packaging was purchased from Takara (632271). HEK 293T cells were obtained from ATCC (Manassas, USA). All cells were cultured in Dulbecco’s Modified Eagle’s Medium (Gibco, USA) with GlutaMAX and 10% fetal bovine serum (Gibco, USA), in a humidified atmosphere containing 5% CO_2_ at 37 °C.

### 2.2 Vectors and Cloning

The Infusion cloning strategy (Takara Bio, USA) was used to construct the plasmids for this study. Detailed information and sequences for primers and vectors are provided in Table S1. All fragments used for cloning were amplified with Q5 Hot Start High-Fidelity 2X Master Mix (New England Biolabs, M0494S) and purified using the gel extraction kit (Qiagen, 28704). Vectors and fragments were co-transformed into XL10-Gold cells (Agilent Technologies, 200314) to generate recombinant plasmids. All constructs were verified through Sanger sequencing.

### 2.3 Recombinant Human Adenovirus

Linearized recombinant human adenovirus plasmids expressing Halo-tagged WT BRCA1 (pAdEasy-RGD-Halo-TEV-BRCA1) and the Y1853ter truncated BRCA1 (pAdEasy-RGD-Halo-TEV-BRCA1-Y1853ter) were transfected into AD-X-293 packaging cells using Lipofectamine, following the manufacturer’s instructions to generate recombinant adenoviruses. High-titer stocks were produced through the stepwise amplification of adenoviruses in AD-X-293 packaging cells. The titration of adenoviral stocks was quantified using the Adeno-X Rapid Titer Kit (Clontech, 632250).

### 2.4 Immunoblot Analysis

Protein was extracted using a modified cell lysis buffer (20 mM Tris-HCl (pH 7.4), 100 mM NaCl, 0.05% Nonidet P-40, 1 mM PMSF, 0.2 mM DTT and 1x protease inhibitor mixture from Sigma-Aldrich P2714). The cell pellets were resuspended well by pipetting, and then incubated on ice for 10 min. Cell lysates were clarified by centrifugation at 13,000 g at 4°C for 10 minutes and transferred to fresh microcentrifuge tubes. Protein was quantified using the Rapid Gold BCA Protein Assay Kit (Thermo Scientific, A53227), separated on 4–12% Tris-glycine gels (Invitrogen, EA0375BOX), and transferred to PVDF membranes. Blocking was done using the Licor blocking buffer (LICORbio, 927-70001), and antibody dilutions were prepared with the Intercept T20 (PBS) Antibody Diluent (LICORbio, 927-75001).

For detecting expression of the fusion constructs, membranes were incubated with anti-HA (Cell Signaling Technology, C29F4, 1:5000) antibody, anti-Flag (Millipore F7425, 1:400), anti-β actin (Abcam ab8227, 1:3x10^5^), anti-Halo (Promega G9211, 1:1000) at room temperature for 2 hrs.

For detecting protein biotinylation, membranes were incubated with streptavidin-IRDye 800 CW conjugates (LICORbio, 926-32230, 1:3000) in Antibody Diluent (LICORbio, 927-75001) at room temperature for 50 minutes. Images were acquired with the LI-COR Odyssey CLx Imaging System.

### 2.5 Immunofluorescence Microscopy

Cells were grown in eight-well chamber slides and treated as indicated. Fixative was 4% paraformaldehyde with 3% sucrose at room temperature for 15 minutes. Permeabilizer was 0.1% Triton X-100 in PBS for 10 minutes at room temperature. Cells were blocked with blocking buffer (Thermo Scientific, 37580) for 1 hour at room temperature. Primary antibodies anti-HA Cell Signaling Technology, C29F4, 1:500) were diluted in diluent buffer (Thermo Scientific, 003118) and incubated on fixed/permeabilized cells overnight at 4°C. Secondary antibody goat anti-rabbit Alexa 594 (Invitrogen/Molecular Probes) (1:300), SNAP probe (Biolabs, S9103S, 1:200), streptavidin Alexa Fluor 555 (Themo Scientific, S32355, 1:1000), and streptavidin Alexa Fluor 488 (Themo Scientific, S32354, 1:1000) were diluted in antibody diluent and incubated at room temperature for 1 hour. Nuclei were counterstained with antifade mounting medium with DAPI (Invitrogen S36938).

In detail, data were acquired with a Nikon TI2-E inverted microscope controlled by NIS-Elements Advanced Research software. The microscope was equipped with a Yokogawa CSU- W1 spinning-disk scanner, a Hamamatsu Fusion BT16bt sCMOS camera. HA fusion proteins, streptavidin Alexa Fluor 555 labeled proteins were excited by a 561-nm laser, and their emission was collected through a 605/52 emission filter. SNAP-Cell 505-Star and streptavidin Alexa Fluor 488 labeled proteins were excited by a 488-nm laser, and emission was collected through a 525/36) emission filter. DAPI imaging was collected with a 455/50 emission filter (exited by a 405-laser line). All imaging was conducted using a 40x water immersion objective lens, beginning with the 561 nm wavelength, followed by 488 nm, and concluding with 405 nm, at a Z-depth of approximately 10 µm. The images presented in the figures were processed using the Z-projection with max intensity in ImageJ.

### 2.6 Proximity Labeling and Affinity Purification of the Biotin-labeled Protein

3xHA-TurboID-BRCA1^36^ or 3xHA-TurboID-BRCA1-Y1853ter and SNAP-Flag-BARD1 co-expressed in HEK 293T by transfection using Lipofectamine 3000 according to the manufacturer’s instructions. 24h after transfection, medium was replaced in each sample with warm biotin-containing complete DMEM medium to initiate labeling (made fresh; final concentration = 500 μM biotin (Sigma-Aldrich, B4501)). Cells were incubated with biotin at 5% CO_2_ at 37°C for the desired time. The labeling reaction was stopped by moving the cells onto the ice and gently washing them five times with ice-cold DPBS. Ice-cold DPBS was used to detach cells via pipetting and collect the cell suspension into microcentrifuge tubes. The cells were pelleted by centrifugation at 300 g at 4°C for 3 minutes and the supernatant was removed.

Whole-cell lysates were generated and the amount of protein in each sample was determined in triplicate as described above. For each sample, lysate was added to 250 μL streptavidin magnetic beads (Thermo Fisher Scientific, 88817), which were washed twice with 1 mL lysis buffer. The beads were incubated with 3 mg protein from each sample and with an additional 1 ml lysis buffer were rotated overnight at 4°C. After enrichment, beads were separated from lysate, which was saved as the unbound fraction, and then washed twice with cell lysis buffer (1 mL, 2 minutes at RT), once with 1 M KCl (1 mL, 2 minutes at RT), once with 0.1 M Na_2_CO_3_ (1 mL, ∼10 s), once with 2 M urea in 10 mM Tris-HCl (pH 8.0) (1 mL, ∼10 s), and twice with cell lysis buffer (1 mL per wash, 2 minutes at RT). After the final wash, the beads were transferred in 1 mL cell lysis buffer to fresh tubes. Protein was eluted from the 5% V beads by boiling them for 10 minutes at 95°C in 6 x SDS sample buffer diluted in cell lysis buffer and supplemented with 2 mM biotin and 20 mM DTT for silver staining assay.

### 2.7 Halo Affinity Purification

AD-X-293 cells were co-infected with recombinant adenoviruses Ad-Halo-TEV-BRCA1 and Ad- SNAP-pp-FLAG-BARD1, or Ad-Halo-TEV-BRCA1-Y1853ter and Ad-SNAP-pp-FLAG-BARD1, each at an MOI of 5, in T300 flasks. Cell pellets were harvested 48 hours post-infection and lysed using the previously described lysis buffer. The lysates were incubated on ice for 10 minutes and then passed through a 26G needle with a 1 mL syringe 10 times. The supernatant was incubated with magnetic HaloTag beads (Promega, G7281) at 4°C overnight with gentle rotation.

After incubation, the unbound supernatant was discarded. The beads were washed five times with a wash buffer containing 0.005% Nonidet P-40 in 1× TBS. Bound proteins were eluted with a buffer containing TEV protease and 1 mM DTT for 2 hrs at room temperature. The eluate was collected and analyzed by SDS-PAGE, followed by silver gel staining. Each sample was prepared in triplicate from independent experiments.

### 2.8 Identification by Label-free Quantitative Protein MS Sample preparation for Halo Affinity Samples

Halo affinity purified samples were reduced by adding 50mM TCEP to a final concentration of 5mM followed by incubation at 55°C for 30 minutes. Cysteines were alkylated by the addition of 375 mM IAA to a final concentration of 10mM followed by incubation at room temperature in the dark for 30 minutes. Ice cold acetone was added at a 1:5 ratio followed by incubation at -20°C overnight. After precipitation, samples were centrifuged at 14,000 *g* at 4°C for 10 minutes to pellet the proteins. The supernatant was removed, and proteins were allowed to air dry on the bench top for 15 minutes. The proteins were resuspended in 50ul of 50mM TEAB pH 8 with 2mM CaCl_2_ and proteins were digested by adding 500ng of trypsin and incubating overnight at 37°C at 500 RPM (Thermomixer, Eppendorf). Digestion reactions were quenched by adding 10% formic acid to a final concentration of 1%. Digested samples were centrifuged at 10,000 g for 10 minutes to remove particulates and the supernatant was transferred to a fresh tube for LC-MS/MS analysis. Peptide concentrations were measured using a Nanodrop spectrophotometer (Thermo Scientific) at 205 nm.

### Sample Preparation for TurboID Samples

TurboID^36^ samples were resuspended in 50mM TEAB. Disulfides were reduced by adding 50mM TCEP to a final concentration of 5mM followed by incubation at 55°C for 30 minutes. Cysteines were alkylated by the addition of 375mM IAA to 10mM followed by incubation at 25°C for 30 minutes in the dark. Proteins were digested by adding 500ng of trypsin (0.1 mg/mL) and incubating overnight at 37°C at 900 RPM. Reactions were quenched by adding 10% formic acid to 1%. Streptavidin beads were removed using a magnetic rack, and samples were transferred to clean tubes.

### LC/MS/MS Detection and Data Analysis

Samples were injected using the Vanquish Neo (Thermo) nano-UPLC onto a C18 trap column (0.3 mm x 5 mm, 5 µm C18) using pressure loading. Peptides were eluted onto the separation column (PepMap™ Neo, 75 µm x 150 mm, 2 µm C18 particle size, Thermo) prior to elution directly to the MS. Briefly, peptides were loaded and washed for 5 minutes at a flow rate of 0.350 µL/minutes at 2% B (mobile phase A: 0.1% formic acid in the water, mobile phase B: 80% ACN, 0.1% formic acid in water). Peptides were eluted over 100 minutes from 2-25% mobile phase B before ramping to 40% B in 20 minutes. The column was washed for 15 minutes at 100% B before re-equilibrating at 2% B for the next injection. The nano-LC was directly interfaced with the Orbitrap Ascend Tribrid MS (Thermo) using a silica emitter (20 µm i.d., 10 cm, CoAnn Technologies) equipped with a high field asymmetric ion mobility spectrometry (FAIMS) source. The data were collected by data dependent acquisition with the intact peptide detected in the Orbitrap at 120,000 resolving power from 375-1500 *m/z*. Peptides from halo purified samples with a charge of +2-7 were selected for fragmentation by higher energy collision dissociation (HCD) at 28% NCE and were detected in the ion trap using a rapid scan rate. Peptides from TurboID samples with a charge of +2-7 were selected for fragmentation by higher energy collision dissociation (HCD) at 28% NCE and were detected in the Orbitrap at 30,000 resolving power. Dynamic exclusion was set to 60s after one instance. The mass list was shared between the FAIMS compensation voltages. FAIMS voltages were set at -45 (1.4 s), -60 (1 s), -75 (0.6 s) CV for a total duty cycle time of 3s. Source ionization was set at 1700 V with the ion transfer tube temperature of 305°C. Raw files were searched against the human protein database downloaded from Uniprot on 05-05-2023 using SEQUEST in Proteome Discoverer 3.0 with methionine oxidation (+15.995 Da) allowed as a variable modification, cysteine carbamidomethylation as a static modification (+57.021 Da), and variable biotin modification (+226.078) allowed on Lys residues for the TurboID samples. Normalization of the samples was performed. Briefly, the raw abundances for each sample were summed, and all samples corrected such that the individual summed samples equaled the maximum summed raw abundance value. Abundances, normalized abundances, abundance ratios, and p-values were exported to Microsoft Excel for further analysis.

### Data availability statement

The mass spectrometry data have been deposited to MassIVE. The accession number for the data reported in this paper is MSV000095782 (Halo) or MSV000095783 (TurboID) Data are also available on ProteomeXchange (PXD055589, PXD055591). Review access can be obtained using the following username and password:

Halo affinity samples:

Username MSV000095782_reviewer Password: JensenChen_24 ftp://MSV000095782@massive.ucsd.edu

TurboID samples

Username: MSV000095783_reviewer Password: JensenChen_24 ftp://MSV000095783@massive.ucsd.edu

### 2.8 Bioinformatic Analysis

#### Data Analysis

We established specific criteria for the proteins to identify the high-confidence interactions from our comparison of WT and mutant proteins in TurboID and Halo experiments. Each protein had to be supported by at least two peptides, detected in at least two replicates, and show a significant log-fold change ratio of 1.4 in absolute value, along with a p-value of less than 0.05 (refer to Supplementary Tables 2 and 3). After applying these filters, 101 proteins met the criteria in our TurboID experiments, and 137 proteins met the criteria in the Halo experiments.

Venn diagrams were generated by Venny 2.1 using the proteins identified from each sample group, excluding the proteins identified in the contaminants protein database(https://bioinfogp.cnb.csic.es/tools/venny/index.html).

All identified protein counts are based on "Accession” counts rather than "Gene Symbol” counts.

### Pathway Analyses

We utilized the Reactome database to assess pathway enrichment in our datasets using ShinyGo in the IDEP application^37^. All query genes were converted to ENSEMBL gene IDs or STRING-db protein IDs. The gene ID mapping and pathway data are primarily sourced from these two databases. Enrichment p-values were calculated using the hypergeometric test. To account for multiple testing, we computed the False Discovery Rate (FDR) using the Benjamini-Hochberg method. The Fold Enrichment of a pathway is defined as the ratio of the percentage of genes in our list to the corresponding percentage in the background genes. Only pathways with two or more genes are considered for the enrichment analysis. Following the analysis, pathways are initially filtered by an FDR cutoff of 0.05. Significant pathways are then ranked based on FDR and Fold Enrichment. We utilize the "Remove redundancy" option to eliminate similar pathways that share 95% of their genes and 50% of the words in their names and represent them with the pathway with the highest significance.

#### 3D modelling studies

To investigate the interaction between the BRCT domain from BRCA1 WT (NCBI Reference Sequence: NP_001394523.1) and the Y1853ter mutant with BARD1 (NP_000456.2), HSPA1L (NP_005518.3), HSP90AB1 (NP_001258898.1), HSPA1B (NP_005337.2), HSPA8 (NP_006588.1), hnRNPA2B1 (NP_002128.1), hnRNPA3 (NP_001317176.1), and hnRNPAB (NP_004490.2), we employed a combination of computational approaches. Using the AlphaFold protein structure prediction system, we generated high-confidence 3D models of the BRCA1- WT and BRCA1-Y1853ter complexes. To analyze protein-protein interactions, we generated contact maps by calculating pairwise atomic distances (<30 Å) between residues in the predicted structures. Residues within a 0.4 Å distance threshold were considered to be in contact. For visualizing these interactions, we used ChimeraX-1.8^38^ to map residue contacts with default settings, excluding intra-residue and intra-molecule contacts. The contact maps, which detail atomic interactions, were then visualized using the Matplotlib library, with rows and columns representing the amino acid residues of BRCA1-WT or BRCA1-Y1853ter and their binding partners.

## 3. Results

### 3.1 Application and validation of the TurboID proximity labeling system for BRCA1 constructs

BRCA1-Y1853ter is currently known as the smallest clinically identified truncation pathogenic mutation. Due to the adenine duplication at position 5558 nt, a premature stop codon is introduced, leading to the truncation of the final 11 amino acids. Among these, there are three amino acids which are conserved within the BRCT-C domain, highlighted in blue (Figure 1A-B) ^35,39^. This nonsense mutation at the BRCT domain leads to truncated protein products that often lose crucial functions, significantly associated with cancer occurrence. We analyzed the variations in the interactome between WT BRCA1 and the BRCA1-Y1853ter mutant to elucidate the specific functions impacted by this minimal truncation.

**Figure 1.**
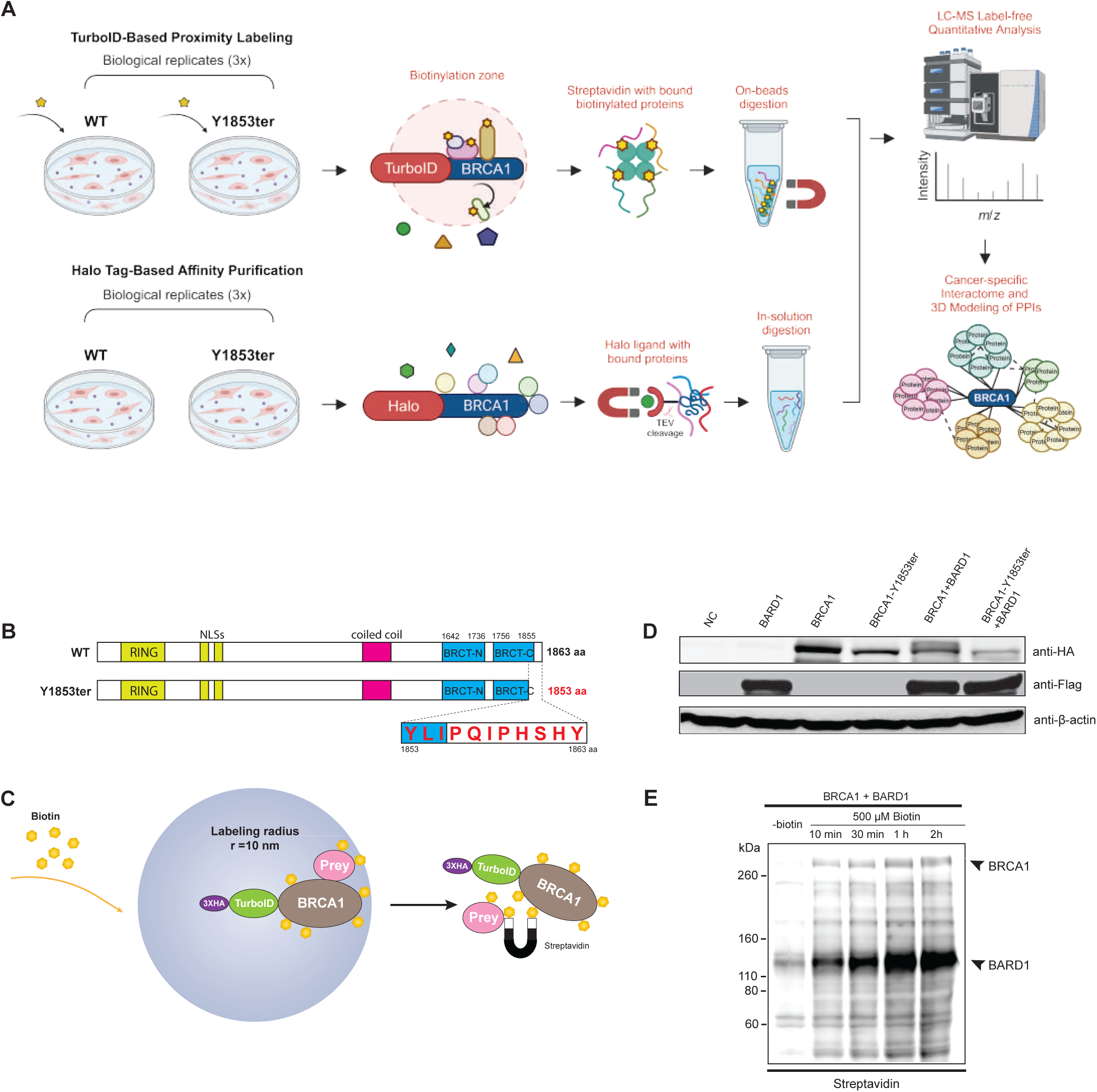
Establishment and optimization of a TurboID-based proximity labeling system for investigating the BRCA1 interactome. **(A)** Workflow for comparative protein interaction analysis using TurboID-based proximity labeling and Halo-Tag-based affinity purification of WT and Y1853ter BRCA1. **(B)** Schematics of BRCA1 and BRCA1-Y1853ter. BRCA1 is a tumor suppressor protein comprising several functional domains including the Really Interesting New Gene (RING) domain for E3 ubiquitin ligase activity, Nuclear Localization Signals (NLSs), and the BRCA1 C-terminus (BRCT) domain involved in transcription activation and DNA repair. BRCA1-Y1853ter lacks the C-terminal 11 amino acids, including three residues within the BRCT-C domain, indicated in red. **(C)** Use of proximity-dependent biotinylation mediated by TurboID to identify BRCA1 interactome. TurboID, a genetically engineered promiscuous biotin ligase derived from a bacterial enzyme, was fused in frame with a 3xHA tag at the N-terminus of BRCA1. This fusion protein utilizes ATP and biotin to catalyze efficient biotinylation of proximal proteins within a 10 nm radius under non-toxic conditions in live cells. Biotinylated proteins were subsequently enriched using streptavidin magnetic beads and identified by mass spectrometry. **(D)** Expression of BRCA1 and BARD1 individual and co-expression analyzed via Western blot (WB). TurboID-fusion proteins were probed using anti-HA antibodies, while BARD1 expression was detected using anti-Flag antibodies. Anti-β-actin antibodies served as a loading control**. (E)** Biotinylation analysis via WB under varying biotin supplementation conditions and time courses (10 min, 30 min, 1 hr, and 2 hrs). Biotinylation levels were assessed using streptavidin-IRDye 800 CW conjugates to probe for biotinylated proteins.

TurboID is a system that utilizes the efficient biotin ligase enzyme that facilitates the study of PPIs by labeling proteins in their native cellular environment and allowing for their subsequent isolation and visualization^36^. We fused 3xHA and TurboID to the WT and mutant BRCA1 N-terminus for expression (Figure 1C). To ensure optimal stoichiometric formation of BRCA1- BARD1 complexes and to facilitate the identification of biologically significant differential interactors, BARD1 was co-expressed in each sample group. Western blot (WB) analysis verified the expression of each fusion protein. As depicted in Figure 1D, HA antibody staining of 293T cell lysates revealed distinct bands corresponding to both WT and mutant BRCA1 at approximately 255 kDa (with BRCA1 at 220 kDa and TurboID at 35 kDa; HA tag size is negligible). This region was absent in empty vector control and BARD1 single-expressed samples, indicating successful expression of the fusion proteins. Similarly, expression of the SNAP-FLAG-BARD1 fusion protein was confirmed using a FLAG antibody at approximately 110 kDa (with BARD1 at 110 kDa and SNAP at 23 kDa; Flag tag size is negligible). The expression level of the mutant BRCA1-Y1853ter, regardless of BARD1 co-expression, was reduced compared to WT BRCA1. This reduction may be associated with instability of the BRCT domain induced by the mutation ^39,40^. Interestingly, during co-expression, cells seem to preferentially maintain BARD1 levels, which remain stable irrespective of BRCA1 co-expression. Conversely, BRCA1 expression is diminished when co-expressed with BARD1. This decrease in BRCA1 levels may be attributed to reduced transfection efficiency associated with the co-expression of BARD1.

The duration of biotin labeling is a critical factor in effectively selecting target proteins. An insufficient labeling period fails to enrich the targets adequately, whereas an excessively prolonged period can lead to nonspecific binding. We conducted a series of time-gradient experiments to determine the optimal labeling duration for BRCA1 bait. As illustrated in Figure 1E, when WT BRCA1 and BARD1 were co-expressed in the WB assay, a band above 110 kDa, presumably corresponding to the BARD1 fusion protein, was detectable even in the absence of exogenous biotin, suggesting that endogenous biotin labeling was sufficient for detecting the expressed BARD1. Additionally, a faint band corresponding to over-expressed BRCA1 was observed (the leftmost lane). After treatment with 500 µM biotin for just 10 minutes, the bands for BARD1 and BRCA1 showed significant enhancement in the whole cell lysate. In contrast, the bands around 60 kDa did not exhibit noticeable changes, indicating the specificity of the intensity increases in the relevant bands. As the labeling duration increased, the intensity of biotin-labeled BRCA1 and BARD1 bands also intensified. However, no significant differences were observed in the smear regions at 10 minutes, 30 minutes, 1 hour, and 2 hours. This indicates that a 10-minute biotin incubation achieves labeling comparable to longer durations, consistent with recommendations in methodological articles on the TurboID system ^36^. Thus, we determined that a 10-minute labeling period was sufficient for capturing BRCA1-interacting proteins and used this duration for subsequent experiments.

### 3.2 TurboID-based quantitative analysis of the effect of BRCA1-Y1853ter mutation on the BRCA1 interactome

We scaled up to 150 cm dishes to prepare samples for purification. After labeling the samples with biotin for 10 minutes, we enriched and purified the biotin-labeled proteins using streptavidin magnetic beads, followed by silver staining to confirm the purification efficiency. We employed samples subjected to biotin labeling without exogenous protein expression as parallel negative controls, as depicted in Figure S1A. In contrast to the samples expressing exogenous proteins, the negative control (NC) group did not exhibit bands corresponding to BRCA1 and BARD1 fusion proteins, indicating specific and successful enrichment and purification in the sample groups. We observed bands at lower than 260 kDa, within the range of 110-160 kDa, and around 50 kDa in all three sample groups. These bands could be attributed to background signal from 500 µM biotin or labeling of substrates by endogenous BRCA1/BARD1 acting as an E3 ubiquitin ligase. Differences in band distribution and intensity between the WT and mutant lanes were detectable on silver-stained gels (Figure S1A Due to the limited volume of samples available, we opted not to perform quantification or normalization of the samples loaded onto the silver-stained gels to preserve material for MS analysis. Consequently, the assessment of differences relies solely on the quantitative and normalized data obtained from MS.

MS data revealed that we identified up to 3,257 proteins in both WT and Y1853ter samples. Among these identified proteins, a total of 3,113 were found in both groups, with 2,589 of these proteins not included in the latest BRCA1 interaction data from BioGRID (Figure 2A, S1C-D, Table S2), indicating that the TurboID method enabled us to identify many previously unknown interacting proteins. Gupta et al. employed the ascorbate peroxidase derivative (APEX) proximity labeling method to characterize the interaction network of WT BRCA1 ^41^. Our analysis identified 1,235 shared hits with their findings (Figure 2B, S1B). Among these identified proteins, BRCA1 had the highest number of peptides (#PSMs = 3263), with BARD1 being one of the most abundant proteins. In the mutant, BRCA1’s abundance was only 32.4% of that in the WT, while BARD1’s abundance was 66.1% of the WT level (Figure S1E, Table S2). The decrease in BRCA1 abundance is consistent with our observations from WB experiments. Although BARD1 expression did not show significant changes in WB analysis, the mutation in our study is located in a non-RING domain, leading us to speculate that the observed decrease in peptide numbers in MS might be due to reduced expression of the mutant. The top ten most abundant proteins are predominantly lower in abundance in the mutant samples compared to the WT samples (Figure S1E, Table S2).

**Figure 2.**
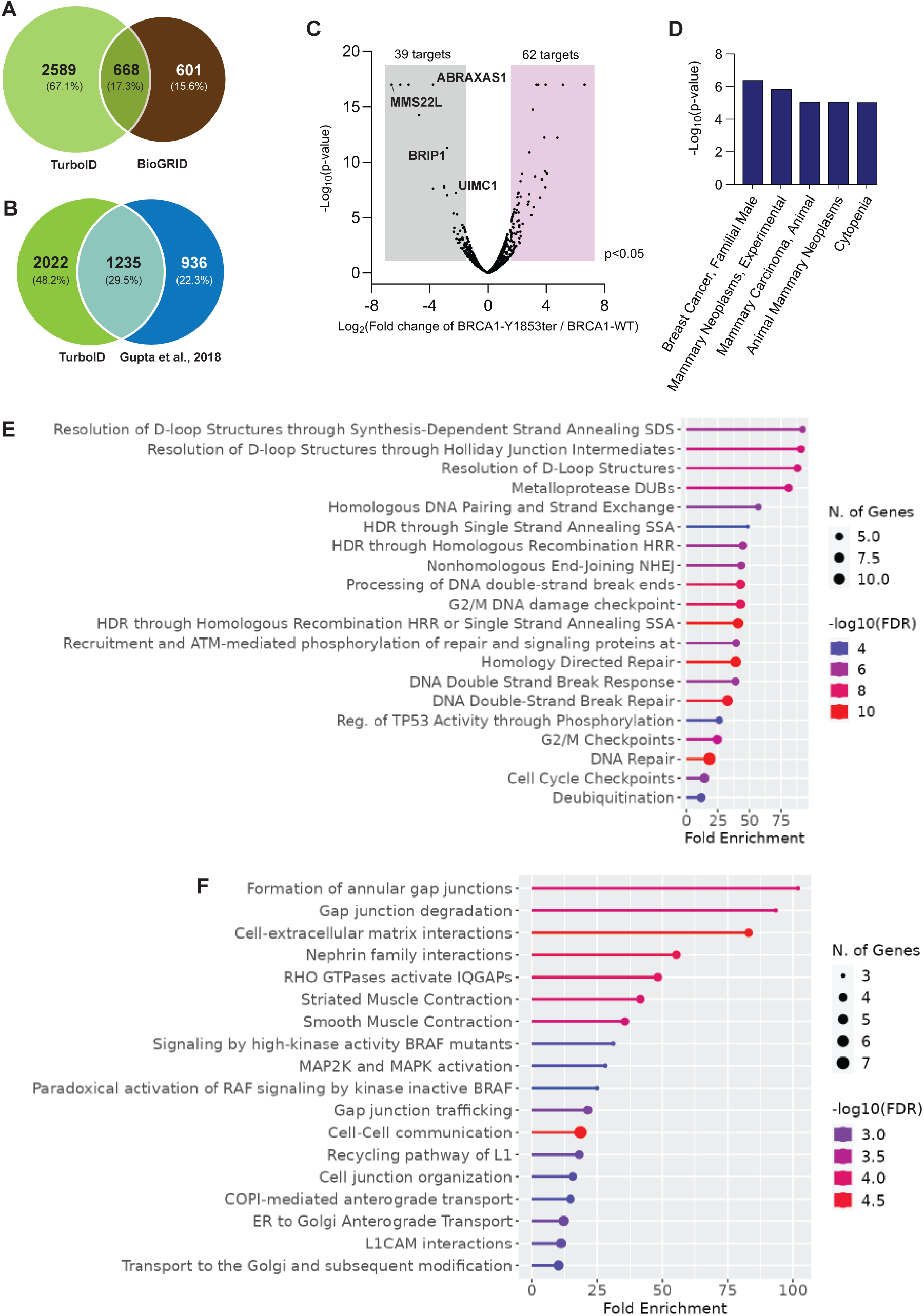
Quantitative analysis of the impact of the Y1853ter mutation on the BRCA1 interactome using TurboID-based proximity labeling. **(A)** The majority (67.1%) of the identified PPIs in this study are not represented in the BioGRID BRCA1 interactome database. **(B)** Venn diagram illustrating the overlap of PPIs identified in this study using TurboID-based proximity method and matching proteins identified through the APEX2-based proximity labeling method in U2OS cells (Gupta et al.^41^). **(C)** Following MS data analysis, TurboID-BRCA1 WT and TurboID-BRCA1-Y1853ter candidates were analyzed and submitted to UniProt using the “Retrieve/ID Mapping” tool. A volcano plot illustrates the log2 ratio of candidates in the comparative analysis between WT and BRCA1-Y1853ter. Enhanced binding with WT BRCA1 is depicted in the grey square (left), while enhanced binding with mutant BRCA1 is shown on the right (blue). Reproducibility across biological replicates was used as an initial filter, selecting proteins that have at least two peptides and at least two replicates in either the WT or mutant samples. This was followed by applying log2 (mut/WT) ratio and log10 P-value cutoffs to further refine the proteome, as indicated by the dashed lines. **(D)** Enrichment analysis based on the DisGeNET database, revealing diseases which are associated with the changes in PPIs. X-axis shows the number of genes involved in each disease. **(E)** PPIs downregulated in the mutant vs. WT BRCA1. **(F)** PPIs upregulated in the mutant vs. WT BRCA1. P-values reflect the statistical significance of enrichment, with lower values indicating a lower likelihood of the result occurring by chance (null hypothesis). False Discovery Rate (FDR) q-values adjust P-values for multiple testing to control type I errors. Fold Enrichment measures the magnitude of enrichment, with higher values indicating stronger enrichment and serving as an important metric of effect size. These results provide crucial insights into the impact of our research.

Further data filtering revealed 101 high-confidence interactions from our WT and mutant comparison. Among these, 39 interactions showed a higher degree with WT BRCA1. In comparison, 62 interactions exhibited a greater change in abundance ratio with the mutant BRCA1-Y1853ter (Log2 cutoff 1.4, p-value < 0.05, and appearing in at least 2 out of 3 biological replicates with at least two peptides) (see Figure 2C, Table S2). The list includes known BRCA1- interacting proteins ABRAXAS1, MMS22L, BRIP1, and UIMC1, which are consistent with previous studies showing a reduction in binding affinity for BRCA1 mutants ^42^. The abundance comparison of the ten most abundant proteins identified in both the WT and mutant datasets are illustrated in Figure S1F.

Since our experiments were conducted in non-cancer cells, we investigated whether the identified interactions might be relevant to cancer. Disease enrichment analysis revealed several categories highly associated with breast cancer, including mammary neoplasms and carcinoma. Additionally, cytopenia, which is strongly associated with Fanconi anemia (FA), suggests an intersection between FA pathways and BRCA1 function (Figure 2D, Table S2) ^43^.

We conducted a comprehensive signaling pathway enrichment analysis to investigate the alterations in biological activities resulting from interactome modifications. Our analysis revealed that the advantageous interactions retained by WT BRCA1, as compared to its mutant form, are associated with multiple critical signaling pathways. These include homologous recombination DNA repair, non-homologous end joining (NHEJ), cell cycle regulation, TP53-mediated regulation, mitochondrial genome maintenance involving D-loop structures, and deubiquitylation processes (Figure 2E, Table S2). Notably, the impairment of DNA repair and cell cycle regulation pathways has been extensively validated as intricately linked to cancer development driven by BRCA1 mutations ^44–47^.

Conversely, we observed that the MAPK signaling pathway activated in BRCA1-Y1853ter mutation, potentially due to the acquisition of interactions that differ from those of WT BRCA1. Our results suggest a potential activation mechanism involving BRAF mutations (Figure 2F, Table S2). In cancer cells, dysregulated activation of the MAPK pathway has been implicated in driving uncontrolled proliferation, evasion of apoptotic mechanisms, and enhanced migratory and invasive properties ^48^. Although high-frequency alterations in MAPK signaling have thus far been reported primarily in low-grade serous ovarian cancer (LGSC) ^49^, and BRCA1/2 mutations are positively correlated with an elevated risk of high-grade serous ovarian cancer (HGSC), our findings propose a potential association between MAPK pathway dysregulation and BRCA1 mutations. Additionally, RHO GTPases play a crucial role in cancer initiation and progression, as evidenced by our enrichment analysis (Figure 2F). In tumor samples from BRCA1-mutant breast cancer patients, alterations in RHO GTPase expression levels were observed, which may correlate with the aggressiveness and metastatic potential of the tumors ^50^

### 3.3 Halo-Based AP-MS reveals functional changes in the BRCA1 interactome

The TurboID-based proximity labeling method identified a large array of targets; however, some of these targets may be artifacts due to labeling by endogenous biotin ligases, leading to potential false positives. The duration of labeling and the 10 nm labeling range further heightened the risk of non-specific labeling. To address these concerns and obtain reliable interactome data, we employed Halo-based AP-MS to compare the interactomes of WT BRCA1 and mutant BRCA1-Y1853ter (Figure 1A, 3A). Unlike traditional affinity tags, the Halo affinity tag provides irreversible binding to ligand-coated magnetic beads, facilitating the capture of transient or weak but specific interactions. This makes it an optimal method for orthogonal comparison with TurboID. While TurboID enriches prey proteins, Halo-based affinity purification relies heavily on the expression levels of the bait protein. We utilized a recombinant adenovirus expression system to ensure comparable prey sample quantities between Halo AP-MS and TurboID methods. A 33 kDa Halo tag, similar in size to TurboID, was fused to the N-terminus of both WT and mutant BRCA1 constructs, and each sample was co-infected with recombinant adenovirus expressing BARD1.

We carried out WB analysis to confirm the expression of all constructs (Figure 3B). Consistent with TurboID findings, the expression level of the mutant BRCA1 fusion protein was lower than that of WT. BARD1 and WT BRCA1 exhibited comparable expression levels in both singular and co-expression conditions; however, mutant BRCA1 showed reduced expression when co-transduced. By purifying and enriching Halo-BRCA1 fusion proteins and their interacting partners using Halo ligand beads, we subsequently released them from the beads with TEV protease *in vitro*. Compared to whole cell lysates (WCL), our data demonstrated effective purification of the samples with silver staining (Figure S2A). The purified samples revealed more distinct bands in the BARD1 co-expression samples than those with BRCA1 alone and endogenous BARD1, with a band above 110 kDa, similar to results obtained with the TurboID method. BRCA1 appeared slightly below 260 kDa, reflecting the successful cleavage of the 33 kDa Halo tag. This result further confirms that the binding affinity between N-terminally tagged BRCA1 and BARD1 was not adversely affected by the 33 kDa Halo tag.

**Figure 3.**
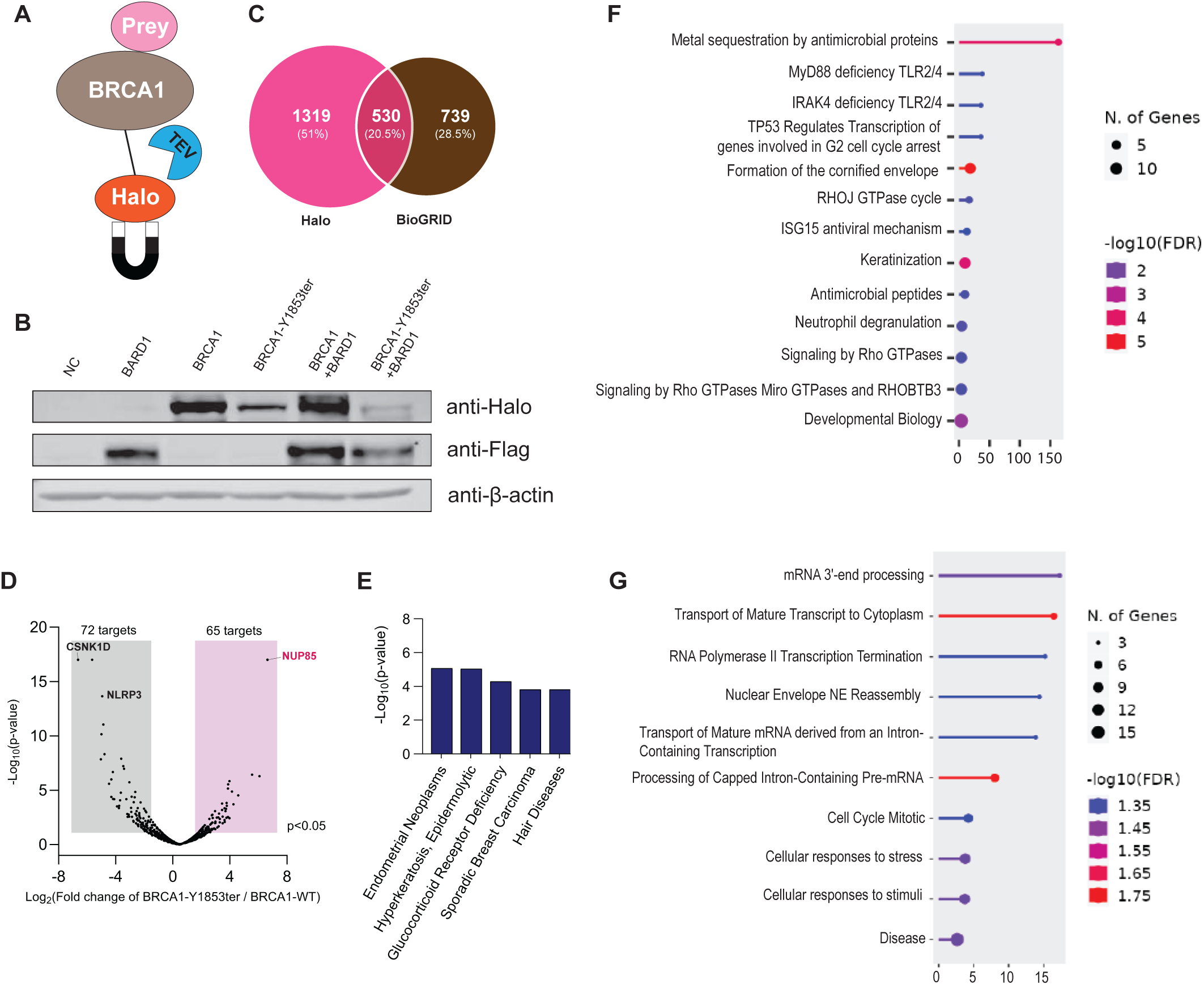
Quantitative analysis of the impact of the Y1853ter mutation on the BRCA1 interactome using Halo-based affinity purification. **(A)** Schematic of the Halo-base APMS method: Halo fusion at the N-terminus of BRCA1 facilitates the purification of BRCA1 and its interactors via TEV protease cleavage in vitro, enabling their release from Halo ligand beads. **(B)** Expression of BRCA1 and BARD1 individual and co-expression analyzed via WB. Halo-fusion proteins were probed using anti-Halo antibodies, while BARD1 expression was detected using anti-Flag antibodies. Anti-β-actin antibodies served as a loading control. **(C)** 28.5% of the identified PPIs in this study are not represented in the BioGRID BRCA1 interactome database. **(D)** Following MS data analysis, Halo-BRCA1 WT and Halo-BRCA1-Y1853ter candidates were analyzed and submitted to UniProt using the “Retrieve/ID Mapping” tool. A volcano plot illustrates the log2 ratio of candidates in the comparative analysis between WT and BRCA1-Y1853ter. Enhanced binding with WT BRCA1 is depicted in the grey square (left), while enhanced binding with mutant BRCA1 is shown on the right (blue). Reproducibility across biological replicates was used as an initial filter, selecting proteins that have at least two peptides and at least two replicates in either the WT or mutant samples. This was followed by applying log2 (mut/WT) ratio and log10 P-value cutoffs to further refine the proteome, as indicated by the dashed lines. **(E)** Enrichment analysis based on the DisGeNET database, revealing diseases which are associated with the changes in PPIs. X-axis shows the number of genes involved in each disease. **(F)** PPIs downregulated in the mutant vs. WT BRCA1. **(G)** PPIs upregulated in the mutant vs. WT BRCA1. P-values reflect the statistical significance of enrichment, with lower values indicating a lower likelihood of the result occurring by chance (null hypothesis). False Discovery Rate (FDR) q-values adjust P-values for multiple testing to control type I errors. Fold Enrichment measures the magnitude of enrichment, with higher values indicating stronger enrichment and serving as an important metric of effect size. These results provide crucial insights into the impact of our research.

MS analysis identified up to 1,849 proteins in both WT and mutant samples, including 1319 novel interactors not listed in the BRCA1 interactome database (BioGRID) (Figure 3C, Table S3). Among these identified proteins, a total of 1647 was found in both groups (Figure S2B-C and Table S3). Consistent with the TurboID approach, BRCA1 exhibited the highest number of peptides (#PSMs = 3,143), with BARD1 among the most abundant interacting proteins. In mutant samples, the normalized abundance of BRCA1 was reduced to 41.8% of the WT levels, while BARD1’s abundance was 46.3% of the WT levels (Figure S2D, Table S3). This observed decrease is consistent with the trends seen in TurboID MS results. Among the ten candidates with the highest abundance, six belong to the heat shock protein (HSP) family, and their abundance is significantly elevated in the mutant samples compared to the WT samples (Figure S2D, Table S3).

Employing the same filtering criteria and analytical methods, we detected 137 high-confidence interactions in comparing WT and mutant samples. Of these, 72 interactions exhibited a higher affinity for WT BRCA1, whereas 65 interactions demonstrated higher abundance with mutant BRCA1-Y1853ter (Log2 cutoff of 1.4, p-value < 0.05, and present in at least 2 of 3 biological replicates with at least two peptides) (see Figure 3D, Table S3). The ten hits exhibiting the most significant changes in up or down ratios are illustrated in Figure S2E. These hits display changes in absolute ratios attributable to their complete absence in either the WT or mutant samples, resulting in a pronounced bias in one group (Figure S2F).

Disease enrichment analysis highlighted an association with sporadic breast cancer, thereby validating the reliability of our data. Endometrial neoplasms, which are tumors of the endometrial lining of the uterus, are known to be correlated with breast cancer. Women harboring *BRCA1* genetic mutations exhibit an elevated risk for both breast and endometrial cancers (Figure 3E, Table S3). Pathway analysis revealed that signaling pathways disrupted by mutant BRCA1 include TP53-mediated G2 cell cycle arrest, RHO GTPase signaling, and Toll-like receptor (TLR) pathways. TLR2 and TLR4 are crucial components of the innate immune system. MyD88 deficiency, a critical adaptor protein in TLR signaling, may lead to alterations in immune responses and potentially influence cancer progression (Figure 3F, Table S3).

Furthermore, the mutant BRCA1 protein displays distinct interaction profiles compared to the WT, which may impact cancer development through modulation of transcription-related activities (Figure 3G, Table S3).

### 3.4 Integration of Interaction Data from TurboID and Halo Orthogonal Screening

We examined the overlap between targets detected by TurboID and Halo methods to accurately discern potential targets. Minimal overlap was observed in the volcano plots from the two approaches, prompting a comparison of the unfiltered interactomes. We found 1,151 overlapping hits, accounting for 29.1% of the total hits identified by both methods (Figure 5A, Table S4). A total of 424 proteins matched the BioGRID BRCA1 interactome database (Figure S3A). The distances between the two methods based on the fold change and show that majority of the proteins are having a small difference (<2 based on the Manhattan distance) using a density plot (Figure S3B-C). Among these 1,151 intersecting preys, 51.8% showed an enhanced affinity for the mutant protein using the TurboID method, while this proportion rose to 66% with the Halo method (Figure 4B, Table S4). Overall, we observed a more significant number of hits with elevated binding affinity to the BRCA1-Y1853ter mutant protein compared to those with stronger affinity for the WT protein. This difference suggests that while there are some variations between the methods, the overall trend in hit variation is highly consistent, reflecting the reliability of both approaches in detecting significant interactions. Heatmap analysis was performed on 390 hits exhibiting increased interaction with the mutant protein and 183 hits showing enhanced interaction with WT BRCA1 (Figure S3D-E, 4C, Table S5). Our data reveal that the Halo method generally detected more pronounced changes in binding affinity than the TurboID method. In the intersection of preys demonstrating consistent trends, the top 10 ratio changes for each methodology are detailed in Figures S3F-I (Table S5).

**Figure 4.**
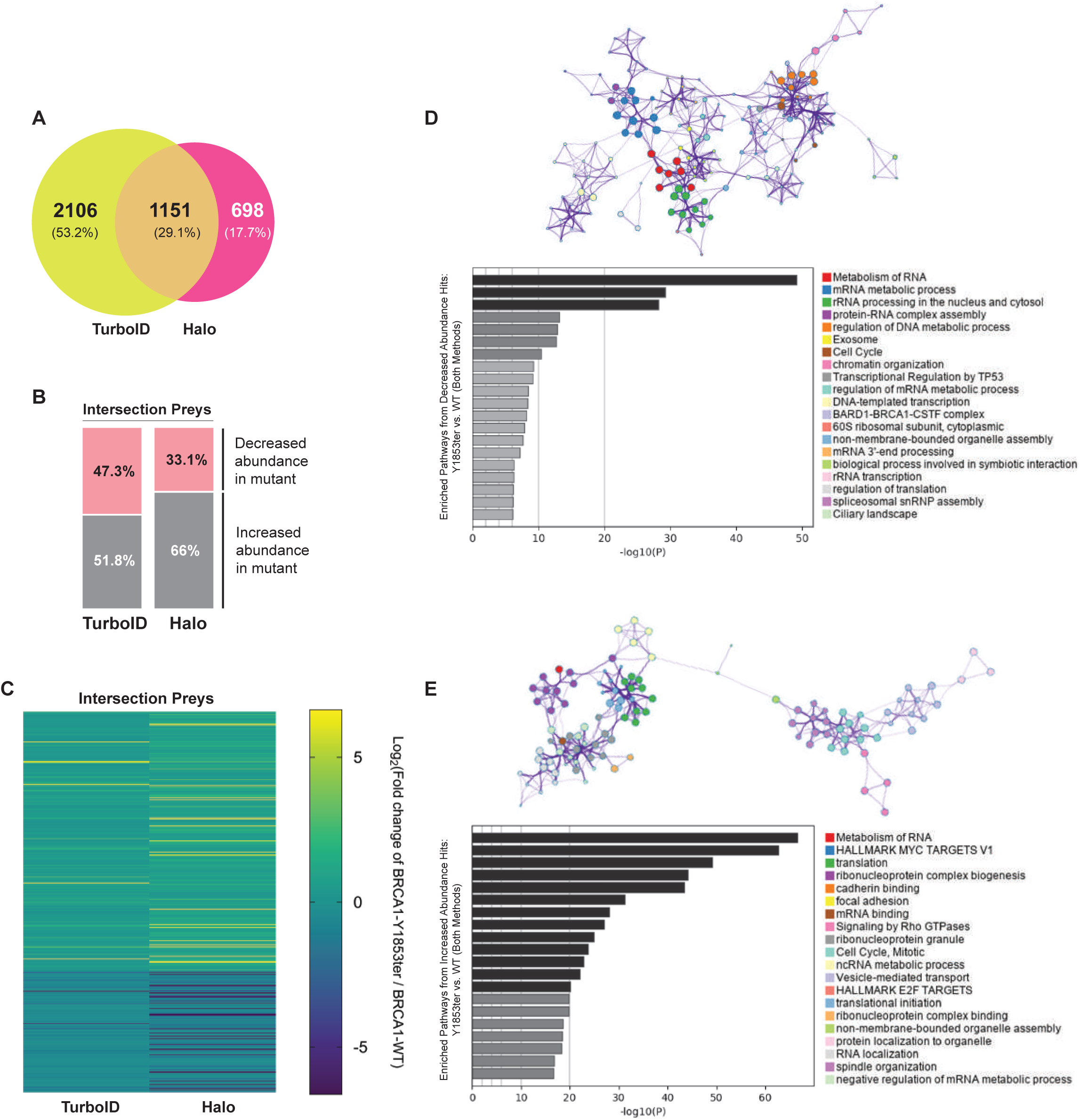
Proteome-scale networks revealed by dual methods: Insights into cancer-specific interactomes and their impact on cellular pathways. **(A)** Overlap among BRCA1 interactions in TurboID and Halo-based networks. **(B)** Intersection interactions exhibit similar trend patterns across two different methods. **(C)** Heatmap depicting quantitative changes in interactions for each intersection component across two methods. **(D)** Metascape performed enrichment analysis on pathways involving intersection downregulated PPIs and assessed the connections between these pathways. **(E)** Metascape performed enrichment analysis on pathways involving intersection upregulated PPIs and assessed the connections between these pathways.

**Figure 5.**
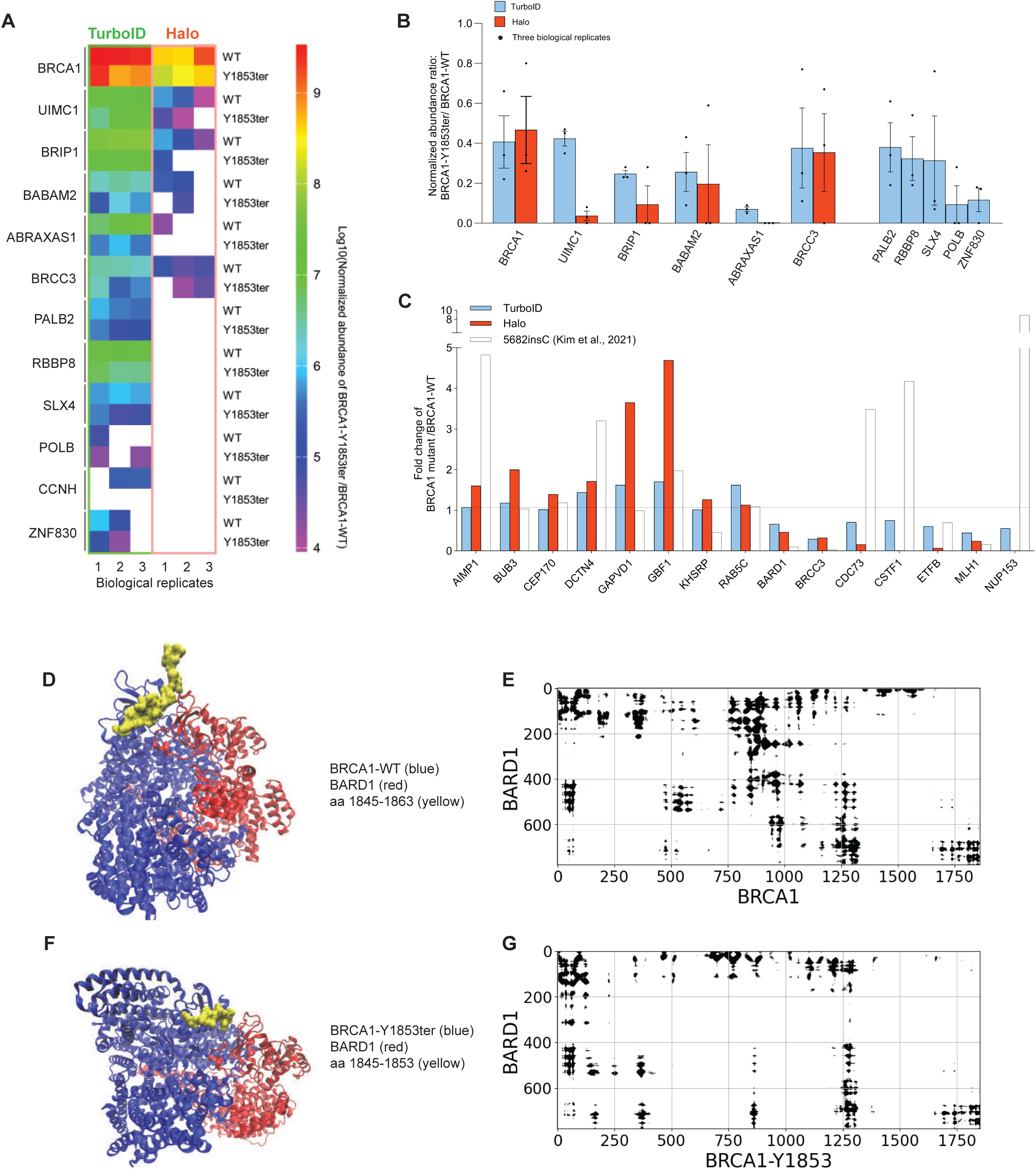
Structural and validation analysis of cancer-specific interactome candidates. **(A)** The heatmap illustrates normalized abundance of individual hits in three biological replicates for each method. **(B)** Ratios of hit abundance variation between WT and mutant samples. Ratio reflecting the abundance variation of each hit between WT and mutant samples, calculated for each method and biological replicate if applicable. **(C)** Bar graph illustrating the common hits between the 183 downregulated and 390 upregulated hits identified in our study and those related to BRCA1-5682insC PPIs from the study by Kim et al., ^25^, along with the ratio changes for each hit across the methods. **(D-G)** AF3-based structural analysis and contact map of WT and mutant BRCA1 interactions with BARD1.

Pathway analysis using Metascape revealed that both downregulated (Figure 4D, Table S5) and upregulated (Figure 4E, Table S5) interaction sets prominently feature RNA metabolism as the most statistically significant biological activity, highlighting the extensive involvement of RNA metabolism in the biological processes associated with BRCA1 mutations. RNA metabolism encompasses a broad spectrum of molecular biology activities and processes, including RNA synthesis, modification, transport, degradation, and translation. This indicates that these processes are differentially regulated due to BRCA1 mutations, some upregulated and others downregulated. For instance, our data indicated alterations in rRNA processing, mRNA processing, protein-RNA complex assembly, translation regulation, and snRNP assembly.

Additionally, downregulated biological activities included DNA metabolic processes, cell cycle TP53-mediated transcription, and chromatin organization. The BARD1-BRCA1-CSTF complex, which plays a role in DNA repair, transcriptional regulation, and mRNA processing, is prominently featured in our list of downregulated activities, and is closely associated with cancer development (Figure 4D, Table S5).

Conversely, our data show that in the context of BRCA1-Y1853ter mutation, cell proliferation-related hallmark genes are upregulated, such as MYC and E2F targets. Upregulated aspects of RNA metabolism include ribonucleoprotein complex biogenesis, translation initiation, ncRNA metabolism, RNA localization, and negative regulation of mRNA metabolism. Rho GTPase and mitotic regulation of the cell cycle are also notable. Furthermore, cytoplasmic-specific processes such as vesicle-mediated transport and spindle organization are among the upregulated activities. These mutation-specific interactions suggest that the biological activities affected by BRCA1 mutation, while appearing relatively independent, may be interconnected, indicating that mutated BRCA1 could contribute to cancer development through a network of interrelated biological processes (Figure 4E, Table S5).

### 3.5 Comprehensive Structural and Comparative Validation of BRCA1 Interactome Candidates

Protein complexes typically exhibit a high degree of conservation in their structural architecture and interaction profiles, particularly among core components, which is crucial for their functional roles in cellular processes. We initially focused on analyzing DNA repair-related complexes, as this is regarded as a crucial function of BRCA1 as a tumor suppressor. Our data reveal that all 12 identified hits exhibited a stable and consistent downregulation of the Y1853ter/WT abundance ratio across three biological replicates for both methods. Notably, the Halo method identified only 6 hits, with CCNH not detected in the Halo method and observed only twice in the WT repeats using the TurboID method (Figure 5A). This strongly underscores the reliability and reproducibility of our data. Furthermore, the downregulation of these hits in the Halo method was generally associated with a greater magnitude of change compared to the TurboID method (Figure 5B).

Although affinity purification methods have been widely employed across various cell systems using different tags and expression systems to investigate BRCA1 interaction proteins, few comprehensive reports analyze the systemic impact of pathogenic BRCA1 mutations on the interactome at the proteomic level. Kim et al. utilized affinity purification to extensively compare the effects of seven BRCA1 clinical mutants on the interactome, including a BRCT truncation mutation, BRCA1-5682insC (an insertion mutation resulting in a 74-amino acid C-terminal frameshift and premature termination, leading to a 35-amino acid truncation). Their study, employing a high PPI score screening system for BRCA1-5682insC, identified 116 specific interacting proteins ^25^. Many of these hits also appeared in the interaction subsets identified by our methods, demonstrating consistent trends. This includes proteins with diminished binding to the BRCA1 mutants, such as BARD1, BRCC3 (BRCA1/BRCA2-containing complex subunit 3), ETFB (Electron Transfer Flavoprotein Beta Subunit), and MLH1 (MutL Homolog 1); as well as proteins with enhanced binding to the BRCA1 mutants, including AIMP1 (Aminoacyl-tRNA Synthetase Complex Interacting Multifunctional Protein 1), BUB3 (BUB3 Mitotic Checkpoint Protein), CEP170 (Centrosomal Protein 170kDa), DCTN4 (Dynactin Subunit 4), GBF1 (Golgi Brefeldin A Resistant 1), and RAB5C (Ras-related Protein Rab-5C) (Figure 5C). These findings underscore the robustness of the intersection subsets identified by our methods.

Utilizing AlphaFold 3 (AF3)^51^, we conducted an in-depth analysis of the BRCA1-BARD1 interaction. The predicted binding patterns revealed subtle alterations in the interaction between mutant BRCA1 and BARD1 relative to the WT complex (Figure 5D, 5F, S4G). Notably, contact mapping indicated that the N-terminal regions of both BRCA1 and BARD1 remained largely conserved, suggesting minimal disruption in these domains. In contrast, a marked reduction in contact frequency was observed between residues 400–600 of BARD1 and residues 750–1000 of BRCA1 ((Figure 5E, 5G). This observed loss of intermolecular contacts is consistent with our mass spectrometry data, which similarly demonstrated a decrease in binding affinity.

### 3.6 Structural Elucidation of Enhanced HSP and hnRNP Families Binding to BRCA1- Y1853ter via AlphaFold 3 and ChimeraX

Moreover, we identified several protein complexes in our analysis of the 390 intersecting hits with elevated affinity for the BRCA1-Y1853ter (Figure S3E, Table S5). Heat shock proteins (HSPs), which are upregulated in response to heat stress and other cellular stress conditions. These molecular chaperones have distinct cytoprotective functions and are crucial for protein folding, repair, and degradation, sustaining cellular integrity and viability. In cancer cells, HSPs are often overexpressed to support the elevated metabolic demands associated with rapid proliferation. HSPs can function either as individual molecules or as part of multi-protein complexes. Current research on BRCA1 has predominantly focused on interactions with HSP70 and HSP90 subunits. It has been reported that BRCA1 stabilizes itself through its interactions with these chaperones (^22,52^). In our screening, we identified eight HSP components, with HSPD1 being the sole component exhibiting reduced affinity for BRCA1-Y1853ter (Table S5). In contrast, the remaining seven components demonstrated enhanced binding to the mutant BRCA1, and this increase was consistently observed across both screening methodologies (Figure 6A). Among these, HSPH1, a member of the HSP105 family, showed a significant fold increase in both methods (1.97-fold in TurboID and 1.93-fold in Halo relative to WT). HSP90AB1 and HSP90AA1 are members of the HSP90 family, while the other four are part of the HSP70 family, with HSPA1B being a well-characterized BRCA1 interactor. STRING analysis reveals these proteins are interrelated, with HSPA8 and HSP90AA1 showing the most significant associations with BRCA1 (Figure 6B).

**Figure 6.**
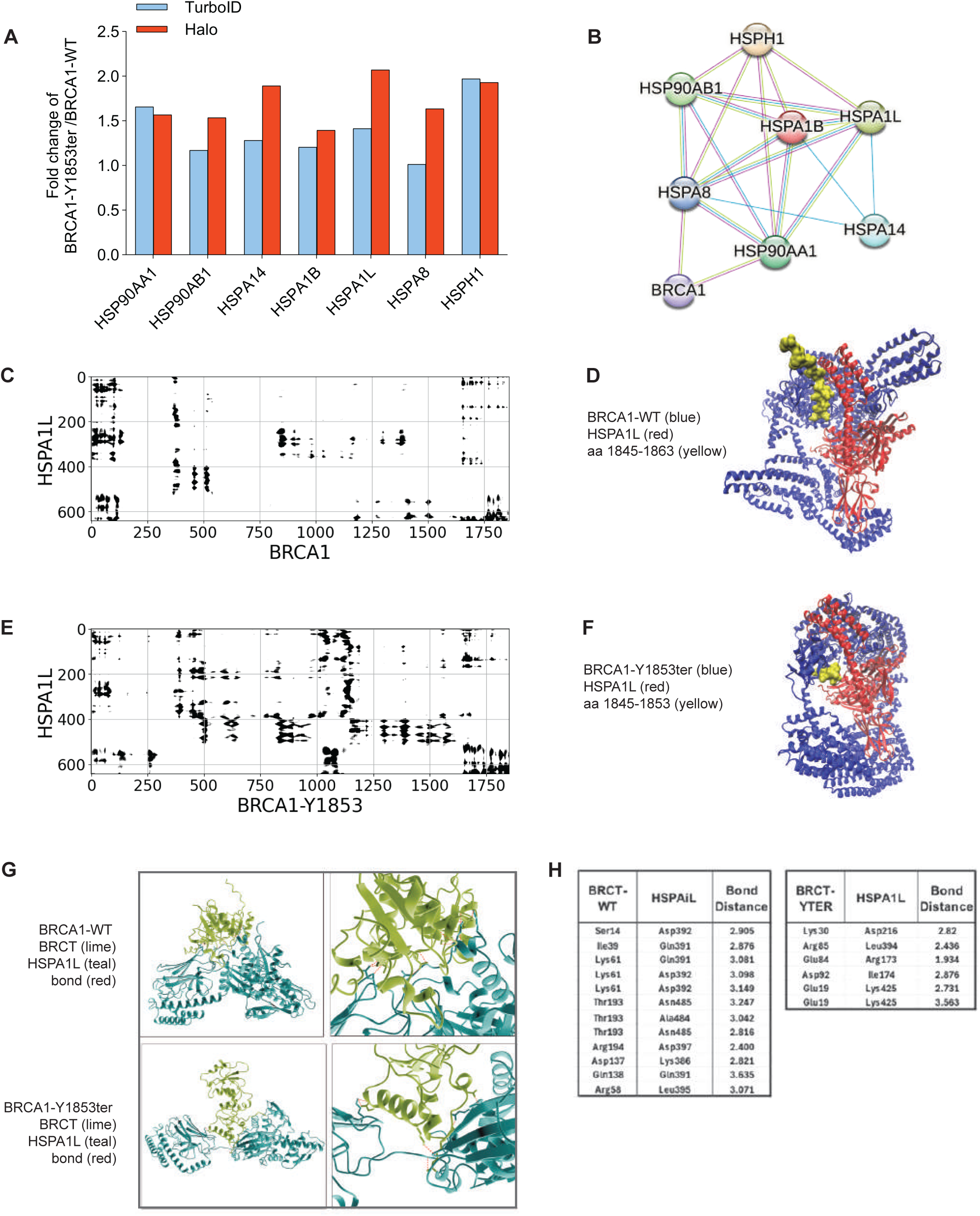
AlphaFold 3 and ChimeraX analysis of enhanced HSPs binding to BRCA1-Y1853ter. **(A)** Variation in the levels of several component proteins identified in the heat shock protein (HSP) complex across two methods. **(B)** STRING highlights interactions and predicts connections between these HSP component proteins and BRCA1. **(C-F)** AF3-based structural analysis and contact map of WT and mutant BRCA1 interactions with HSPA1L. **(G-H)** ChimeraX-based structural analysis and contact map of WT and mutant BRCA1 interactions with HSPA1L.

AF3 structural predictions revealed significant conformational changes in the binding mode of HSPA1L with mutant BRCA1 compared to WT BRCA1 (Figure 6D, 6F). The contact map further demonstrated enhanced binding at the C-terminus of BRCA1, particularly from approximately amino acid 750 to the C-terminus. Notably, the BRCT-N domain of BRCA1-Y1853ter exhibited a pronounced increase in binding affinity with HSPA1L (Figure 6C, 6E). These findings suggest that mutant BRCA1 engages in more dynamic and stabilized interactions with HSPA1L.

Hsp90AB1 exhibited a more substantial shift in binding in the contact map relative to HSPA1B, while HSPA8 showed a notable decrease in binding to mutant BRCA1 at the N-terminus compared to WT (Figure S4A-F). Structural superimposition of the complexes, using the unaltered binding partner (either BRCA1 or BRCA1-truncated) as a reference, revealed distinct differences in the interaction modes of BRCA1 and BRCA1-Y1853ter with their respective partners (Figure S4G-J). In most cases, the truncated form of BRCA1 exhibited altered interaction profiles with the analyzed candidates. However, this effect was less pronounced for BARD1, which is consistent with the predictions from the contact map analysis (Figure 5E, 5G). The superimposed structural models clearly demonstrate that BRCA1-Y1853ter interacts with its partners differently from WT BRCA1, even at regions distant from the truncation site.

The crystal structure of full-length BRCA1 has not yet been resolved due to the large size of the molecule, although structural data for the BRCT domain are available. To further investigate how the deletion of the 11 C-terminal amino acids affects the structure and function of the BRCT domain, we performed an in-depth analysis of the BRCT-HSPA1L interaction. Our ChimeraX^53^ data revealed substantial alterations in the binding mode, although the BRCT domain itself is minimally affected by the truncation mutation (Figure 6G, S5A). The analysis of key amino acid residues showed significant changes in bond distances, which further corroborates the enhanced binding between HSPA1L and mutant BRCA1 (Figure 6H).

Notably, the heterogeneous nuclear ribonucleoprotein (HNRNP) complexes comprised 12 components within this intersection. Of these, 11 components demonstrated a heightened affinity for the mutant BRCA1 across both methodologies, while HNRNPR was the only component showing diminished binding (Figure 7A, Table S5). HNRNPs generally operate as multi-protein complexes, binding to nuclear RNA to facilitate various aspects of RNA metabolism, including processing, transport, and regulation. These complexes are integral to the modulation of RNA splicing, nuclear export, stability, and post-transcriptional modifications. We assessed the variation of these 11 HNRNP components across the two methods. Distinctly, HNRNPH2 exhibited considerable variability, with a 7.78-fold increase in the Halo method compared to WT, and a 1.03-fold increase in the TurboID method. In contrast, the other 10 components showed relatively consistent fold changes. STRING analysis further elucidated that these subunits are involved in known interactions, with HNRNPL, HNRNPH2, HNRNPDL, HNRNPC, and HNRNPA2B1 being established BRCA1 interactors ^54,55^. Moreover, STRING analysis suggested that HNRNPAB, HNRNPA2B1, and HNRNPD may interact more directly with BRCA1 (Figure 7B).

**Figure 7.**
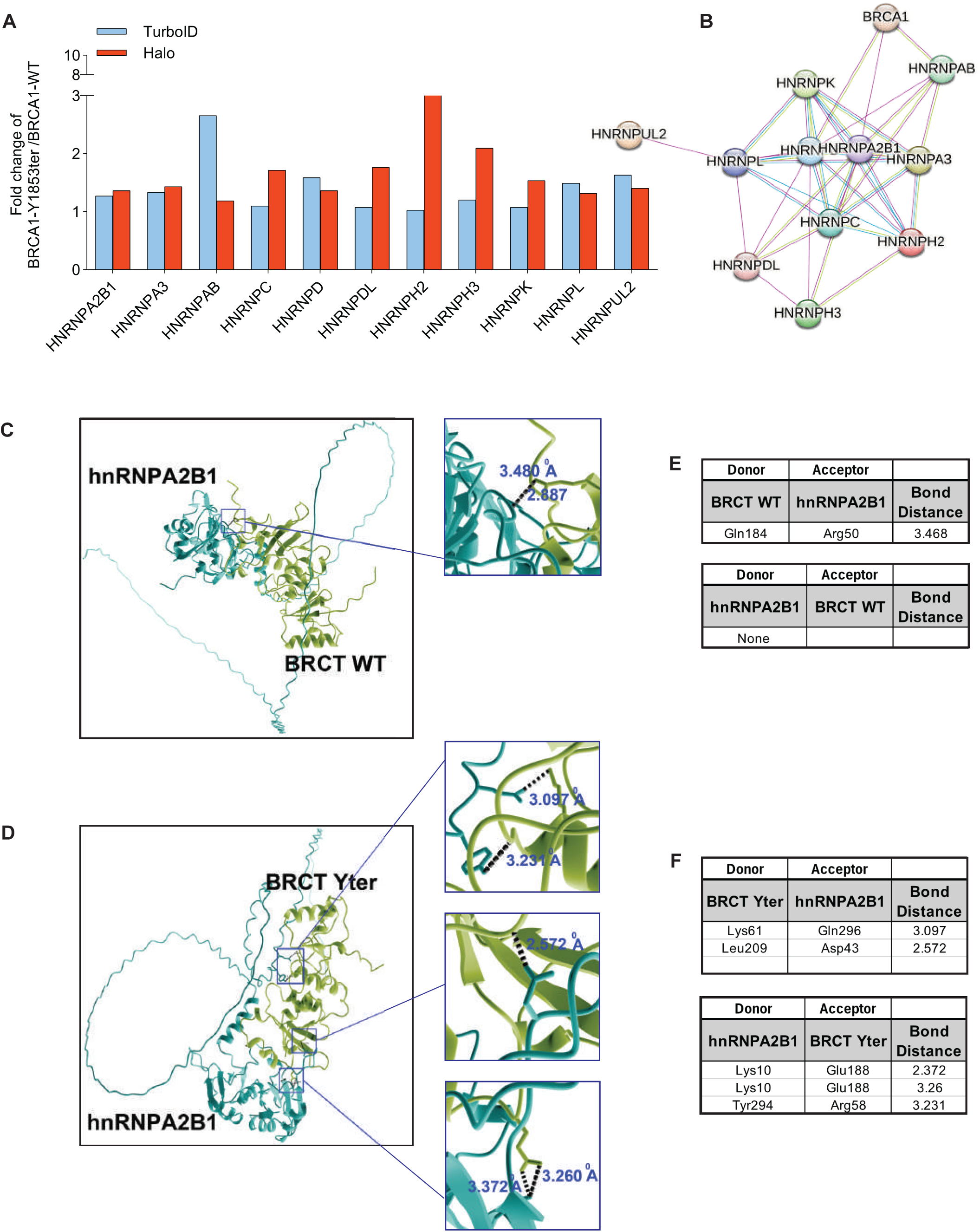
AlphaFold 3 and ChimeraX analysis of enhanced hnRNPs binding to BRCA1-Y1853ter. **(A)** Variation in the levels of several component proteins identified in the heterogeneous nuclear ribonucleoprotein (HNRNP) complex across two methods. **(B)** STRING highlights interactions and predicts connections between these HNRNP component proteins and BRCA1. **(C-F)** ChimeraX-based structural analysis of WT and mutant BRCA1 interactions with hnRNPA2B1.

HNRNPA2B1 has been reported to be significantly upregulated in several subtypes of breast cancer (BRCA) tissues^56–58^. Our ChimeraX analysis reveals that the mutation induces alterations in the BRCT binding mode with HNRNPA2B1, particularly affecting key amino acid residues involved in high-affinity interactions (Figure 7C-D). This change is especially pronounced when HNRNPA2B1 is modeled as the donor (Figure 7E-F). Furthermore, we applied the same approach to analyze other hnRNP family members, and, consistent with our quantitative mass spectrometry data, these proteins demonstrated enhanced binding affinities with BRCA1- Y1853ter (Figure S5D-E).

We also identified three components involved in nuclear export: XPO1, XPO5, and XPOT. These components consistently demonstrated enhanced binding affinity for BRCA1-Y1853ter, and notably, none were present in the down binding group (Figure S5B, Table S5). XPO1, or CRM1, is a key nuclear export receptor that mediates the translocation of proteins and RNA molecules from the nucleus to the cytoplasm by recognizing NES on the cargo. XPO5 is primarily responsible for the transport of mRNA and snRNA, facilitating their roles in cellular translation and splicing. XPOT specializes in the export of pre-miRNA and tRNA, which is essential for RNA processing and protein synthesis. BRCA1’s translocation from the nucleus to the cytoplasm via XPO1 is critical for its functional regulation within the nucleus ^59^. Alterations in BRCA1, such as mutations, can disrupt its interaction with XPO1, potentially leading to aberrant intracellular distribution and impaired functionality in DNA repair ^29,60^. STRING analysis further highlights the interactions among these transport proteins (Figure S5C). These results suggest that mutant BRCA1 may profoundly impact cellular processes by altering its subcellular localization and modifying relevant cargo transport mechanisms.

### 3.7 Signal Pathways of method-specific Interactions and their implications

The observation of distinct interaction remodeling patterns induced by BRCA1-Y1853ter across the two methodologies is to be expected. Notably, the 30% overlap in identified interactions underscores the substantial variability arising from the techniques employed. These methodological discrepancies expand the network coverage, facilitating a more comprehensive understanding of the BRCA1 interactome alterations and providing insights into the advantages and limitations of each approach. Consequently, we conducted pathway analyses on the interactions unique to each method, categorized by those with higher binding to either the WT or BRCA1-Y1853ter.

In the TurboID approach, interactions exhibiting increased binding to the WT BRCA1 are significantly associated with pathways including DNA replication and repair, RNA polymerase activity, and the spliceosome—processes predominantly occurring within the nucleus.

Additionally, pathways such as ribosome biogenesis, citric acid cycle, and amino acid biosynthesis, which occur in the cytoplasm and mitochondria, respectively, were also prominently associated. Noteworthy are the pathways related to nuclear-cytoplasmic transport and cell cycle regulation (Figure S6A, Table S4, S6). These findings indicate that the mutant BRCA1 is likely impaired in these functional areas, particularly in DNA replication, repair, and transcription, aligning with established research ^61^. Conversely, interactions with elevated binding to the mutant BRCA1 are significantly linked to pathways involved in translation in the cytoplasm, Nonsense-Mediated Decay (NMD), selenoamino acid metabolism in mitochondria, and RNA metabolism (Figure S6B, Table S4, S6).

In the Halo approach, interactions with elevated binding to the WT BRCA1 are associated with pathways such as programmed cell death and apoptosis, which are operational throughout the cell. Additionally, pathways including antigen processing, ubiquitination, proteasome degradation, metal sequestration by antimicrobial proteins, neutrophil degranulation, and the formation of the condensed envelope—which occurs in the cytoplasm and inner nuclear membrane—were also significantly represented (Figure S6C, Table S4, S6). These associations suggest that mutant BRCA1 may be compromised in maintaining cellular homeostasis and regulating apoptosis. Conversely, interactions with increased binding to the mutant BRCA1 are notably linked to pathways such as programmed cell death, apoptosis, ubiquitin-dependent degradation of cyclin D, nucleotide metabolism, neutrophil degranulation, polyamine metabolism, assembly of the pre-replicative complex, and removal of Orc1 from chromatin (Figure S6D, Table S4, S6). Orc1, a crucial factor for DNA replication initiation, suggests that BRCA1- Y1853ter may influence DNA replication and cell cycle regulation through modulation of Orc1 removal from chromatin.

We conclude that within the same cellular genetic context, both techniques likely capture important, but distinct alterations in the BRCA1 interaction network which should be confirmed by traditional methods. While the shared interactions identified by both methods may overlook certain valuable interactions, they effectively reduce the inherent false positives of each method, capitalizing on their strengths to capture true and stable interaction changes. This approach significantly enhances our understanding of the fundamental cellular processes remodeled by BRCA1-Y1853ter.

### 3.8 Remodeling of the interactome across multiple cellular compartments

Clinical studies have documented that mutations associated with cancer can cause aberrant cytoplasmic localization of BRCA1, with mutations in the BRCT domain, such as M1775R, 5382insC, and the Y1853ter mutation examined in this study, being identified as primary drivers of this mislocalization ^29,62–64^. Nevertheless, the precise relationship between BRCA1’s subcellular localization and its role in carcinogenesis and tumor progression remains unresolved. This uncertainty is attributable to the impact of antibody specificity and tumor heterogeneity on experimental outcomes, as well as the complexity of BRCA1’s regulated nucleocytoplasmic transport, which is modulated by its interactions with various proteins ^65–68^. To elucidate whether the high-affinity interactions observed with BRCA1-Y1853ter are involved in pathways beyond the cytoplasm, we employed the TurboID technique which utilizes streptavidin conjugates to visualize the subcellular localization and distribution of biotin-labeled preys.

We conducted a time-course immunofluorescence experiment to mitigate non-specific localization arising from extended labeling times. As depicted in Figure 8A, the absence of exogenous biotin resulted in negligible fluorescence signal, demonstrating the low background noise associated with streptavidin dyes. Consistent with established literature, WT BRCA1 predominantly localizes to the nucleus. In contrast, our findings for BRCA1-Y1853ter, shows a pronounced cytoplasmic localization aligning with previous observations in HCC1937, T47D, and MCF-7 cells ^29,60^. We compared biotin labeling for 10 minutes versus 2 hours at a concentration of 500 µM biotin. Both time points yielded clear labeling of interactors, with WT BRCA1 exhibiting consistent nuclear co-localization of preys; however, a slight increase in cytoplasmic distribution was noted in the 2-hour samples. For mutant BRCA1, co-localization of preys was predominantly observed in the cytoplasm, with some cells showing a more extensive distribution. Compared to the 10-minute labeling, the 2-hour labeling resulted in a higher proportion of whole cell distribution of interactors, likely due to staining saturation from the prolonged labeling. Consequently, we determined that a 10-minute labeling duration is optimal for our subsequent experiments.

**Figure 8.**
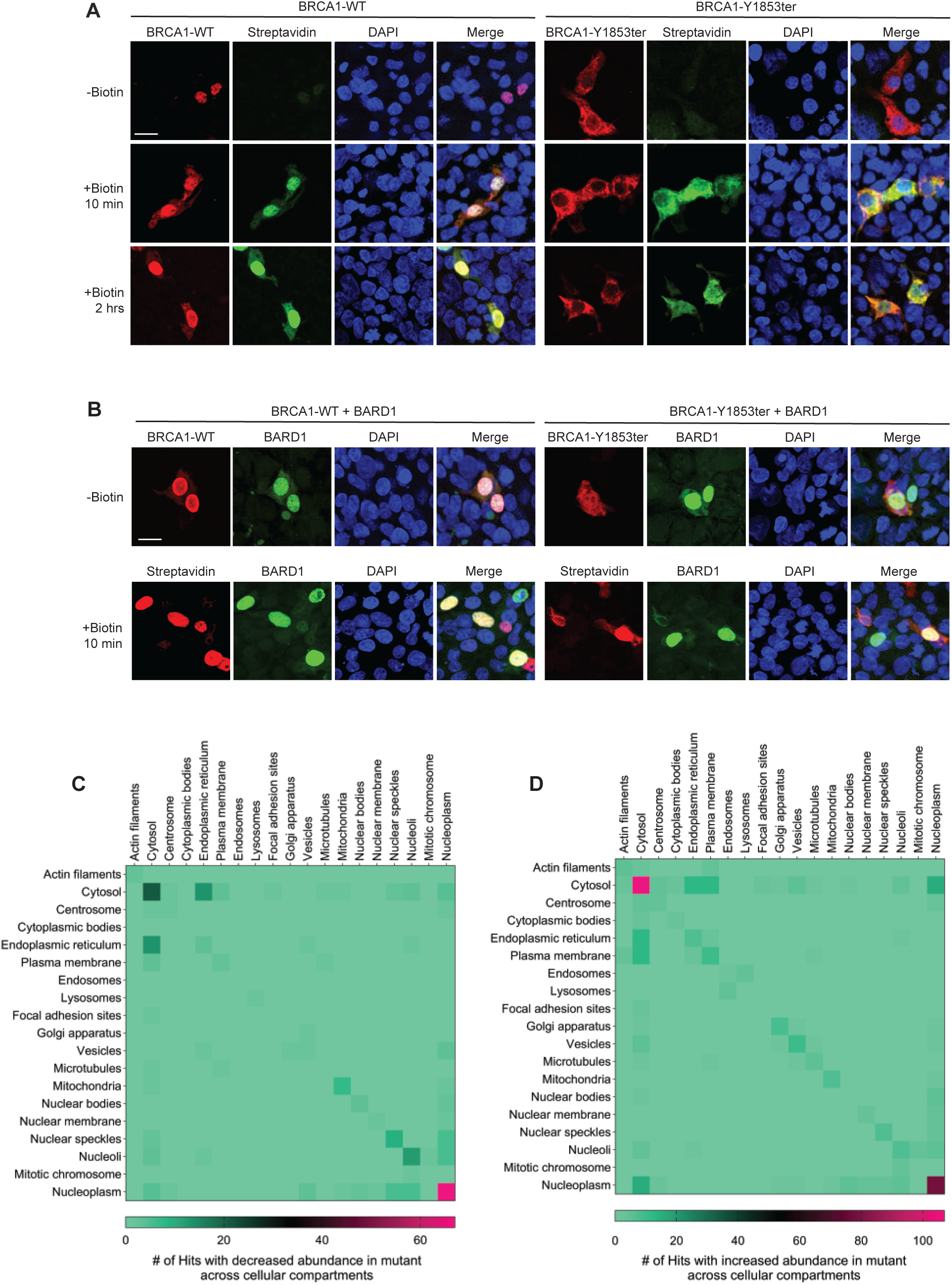
BRCA1-Y1853ter, in conjunction with BARD1, alter interaction networks by causing the loss and gain of PPIs across cellular compartments. **(A)** Visualization of BRCA1 and its binding interactors in 293T cells in different time courses. 293T cells transiently expressing HA-tagged TurboID-BRCA1 and TurboID-BRCA1-Y1853ter were fixed following biotin labeling for indicated times. Anti-HA antibodies were used to visualize fusion protein localization (red). Fluorescently conjugated streptavidin was employed to detect biotinylation following no biotin supplementation (DMSO) or after addition of 500 μM biotin for 10 minutes or 2 hours (green). DAPI staining marked the nucleus (blue). Scale bar: 10 µm. **(B)** Visualization of BRCA1, BARD1, and its binding interactors in 293T cells in the presence of BARD1. 293T cells transiently co-transfected with HA-tagged TurboID-BRCA1 and BARD1 or TurboID-BRCA1-Y1853ter and BARD1 were analyzed similarly. Anti-HA antibodies visualized fusion protein localization (red), while fluorescently conjugated SNAP ligand detected BARD1 distribution. Fluorescently conjugated streptavidin was used to detect biotinylation following no biotin supplementation (DMSO) or after the addition of 500 μM biotin for 10 minutes (green). DAPI staining marked the nucleus (blue). Scale bar: 10 µm. **(C)** Histograms showing the cellular distribution of individual candidates with decreased binding to BRCA1-Y1853ter, and **(D)** those with enhanced binding, indicating the number of corresponding hits in specific cellular compartments.

Studies have demonstrated that BRCA1 harbors multiple nuclear localization signals (NLS) and nuclear export signals (NES), which collectively facilitate its dynamic shuttling between the nucleus and cytoplasm throughout the cell cycle ^69^. Beyond its intrinsic localization signals, BRCA1’s subcellular distribution is also influenced by interactions with various cellular partners. Notably, BRCA1 forms a heterodimer with BARD1, which mediates BRCA1’s nuclear import via BARD1’s NLS. The complex’s NES are masked, allowing the dimer to remain within the nucleus to execute its functions and suppress apoptosis ^32,66,70,71^. BRCA1 interacts with gamma-tubulin in the cytoplasm to modulate centrosome structure and function. Additionally, BRCA1 has been reported to localize to mitochondrial and endoplasmic reticulum compartments ^72^. Prior investigations have shown that reducing BARD1 levels via siRNA or expressing dominant negative BRCA1 N-terminal peptides, which disrupt BRCA1/BARD1 interaction, leads to the loss of BRCA1 from the nucleus ^32^. To further explore this, we examined the effect of co-expressing BARD1 on the localization of BRCA1 and its interactors (Figure 8B). In the presence of BARD1, WT BRCA1 and its interactors and BARD1 itself demonstrated robust co-localization within the nucleus, with only minimal co-localization observed in the cytoplasm. Conversely, for BRCA1-Y1853ter, co-expression with BARD1 enhanced its nuclear localization, similar to previously reported BRCT domain mutants ^29,73^, with a predominant whole cell distribution observed. Furthermore, co-expression with BARD1 led to a significant shift in the localization of BRCA1-Y1853ter interactors from a predominantly cytoplasmic to a more pronounced nuclear co-localization, with an increased presence in the nucleoplasm compared to WT BRCA1 interactors (Figure 8A and the top row of Figure 8B). Notably, the mutant interactors also displayed enhanced cytosolic distribution relative to their WT counterparts (The bottom row of Figure 8B). The localization of BARD1 remained predominantly within the nucleoplasm, unaffected by co-expression with either WT or mutant BRCA1 (Figure 8B).

Utilizing data from The Human Protein Atlas, we assessed the subcellular localization of 183 WT BRCA1 high-binding interactors and 390 high-binding BRCA1-Y1853ter interactors. Our analysis reveals that many of these interactors, akin to BRCA1, exhibit shuttling properties between the nucleus and cytoplasm, demonstrating diverse localization patterns across various cellular compartments. As depicted in Figure 8C, among the WT high-binding interactors, 67 were exclusively nuclear (the largest proportion at 36.6%), 21 were exclusively cytosolic (the second largest proportion at 11.5%), followed by endoplasmic reticulum/cytosol dual localization (7.1%), nucleoli (6.6%), and nuclear speckles (4.9%). Other less frequent subcellular compartments were primarily within the nucleus, including mitotic chromosomes and nuclear bodies (Figure 8C, Table S7).

Conversely, for mutant high-binding interactors, 107 were exclusively cytosol (the predominant category at 27.4%), 78 were exclusively nuclear (the second most prevalent at 20%), followed by nuclear/cytosol dual-localized proteins (3.8%), endoplasmic reticulum/cytosol dual localization (3.1%), and plasma membrane/cytosol dual localization (3.1%). Other less common compartments included cytoplasmic structures such as the endoplasmic reticulum, plasma membrane, vesicles, microtubules, and mitochondria, as well as nuclear compartments including the nuclear membrane, nuclear speckles, and nucleoli (Figure 8D, Table S7). In summary, these findings reveal that, even with BARD1 co-expression, BRCA1-Y1853ter displays distinct localization patterns compared to WT BRCA1. This suggests that the mutant BRCA1 may form unique interactions in the cytoplasm, potentially facilitating gain-of-functions. Additionally, mutations in the BRCT domain of BRCA1-Y1853ter lead to the loss of certain nuclear binding partners but enable new interactions with nuclear, nuclear/cytosol dual-localized proteins, thereby may also acquire specialized activities within the nucleus.

## 4. Discussion

Most tumor suppressors function primarily as proteins, interacting with specific partner proteins to establish intricate networks. These networks play a crucial role in finely regulating relevant biological pathways, thus preventing the proliferation of abnormal cells and the onset of cancer. Mutations in tumor suppressors disrupt these interaction networks, leading to loss, partial loss, or novel formation of interactions. Quantitative analysis of PPI networks can elucidate the subcellular localization of mutant proteins, alterations in protein complexes, and predict their biological implications. This provides a foundational understanding for linking genotype to phenotype, unraveling complex mechanisms of cancer development, and identifying potential therapeutic targets.

MS-based affinity purification and proximity labeling methods are essential techniques for comprehensive proteomic analysis. Affinity purification allows for capturing interaction partners with high specificity and stability due to the high purity of the bait protein, making it easier to identify robust and specific interactions. In contrast, proximity labeling tags proteins within the vicinity of the target protein under physiological conditions, capturing a broad spectrum of interactions, including transient and weak associations. Gupta et al. first applied APEX for proximity labeling of WT BRCA1 interactomes ^41^.Our study introduces TurboID, an advanced biotinylation enzyme that surpasses APEX in labeling efficiency, background signal reduction, and stability in live cells. The synergistic application of these techniques reveals significant advantages: it greatly extends the PPI repertoire, provides a more comprehensive quantitative comparison of the interactome, and facilitates the systematic dissection of complex intracellular networks and their functions. Additionally, it aids in refining interaction zones to identify more accurate and stable protein interactions.

HEK293T cells, characterized by their rapid growth and high expression of exogenous proteins, are ideal for Halo-tag-based affinity purification mass spectrometry and TurboID-based proximity labeling. Although HEK293T is not a cancer cell line, Huttlin et al., reported that the core complexes identified in HEK293T cells show high conservation with those in colon cancer derived HCT116 cells ^74^. Using these methods in the same cellular system, we compared the effects of WT BRCA1 and the clinically relevant minimal truncation mutant BRCA1-Y1853ter on the protein interaction network. Our findings highlight the complex alterations in signaling pathways resulting from the loss of just 11 C-terminal amino acids in BRCA1, which are closely linked to cancer development. This study identifies core complexes and essential proteins implicated in carcinogenesis, providing novel insights into the molecular mechanisms underlying tumorigenesis.

Additionally, this study reveals the impact of technical variability on experimental results, highlighting the advantages and limitations of each method. We used similar cell quantities for both methods in subsequent purification experiments. The Halo affinity purification method relies on the enrichment of the target protein, meaning the target’s abundance directly influences the prey library’s size. In contrast, TurboID labels proteins based on their proximity to the bait, which is less dependent on the bait protein’s expression levels and is advantageous for detecting transient interactions, weak interactions, and membrane-associated targets often overlooked by APMS. Consequently, TurboID identified 1.73 times more preys compared to Halo, a similar observation to the comparative results between the APEX method and APMS ^41^. When applying consistent filters, including biological replicate reproducibility and statistical significance, the number of hits identified by both methods was roughly equivalent in volcano plots. This suggests that TurboID-based prey screening exhibits variability in biological repeats, likely due to the rapid labeling and sensitivity of TurboID affecting specificity and completeness, indicating a need for more precise optimization of biotin concentration and labeling duration.

Moreover, subcellular compartmentalization is crucial for cellular physiological processes. The TurboID method enabled a more explicit characterization of the impairment in nuclear functions such as DNA replication and repair caused by the BRCA1-Y1853ter. Conversely, Halo affinity purification, which involves cell lysis, may lead to the mixing of proteins from different compartments, increasing the likelihood of non-physiological interactions. This can elevate the chances of cytoplasmic-mutant protein interactions with nuclear proteins, resulting in false-positive nuclear interactions and failing to reflect the impairment of DNA repair pathways accurately.

BRCA1-BRCT mutations disrupt protein folding and stability, mislocalizing many mutant proteins to the cytoplasm. Our interactome analysis indicates that the interactions of mutant BRCA1 in the cytoplasm extend beyond protein folding, stability, and degradation, including RNA transport, translation, vesicle-mediated trafficking, and spindle organization. Furthermore, we identified numerous high-affinity binding partners for mutant BRCA1, many of which are nucleoplasmic proteins, particularly in the presence of co-expressed BARD1. Despite treatment with the proteasome inhibitor MG132, the cytoplasmic intensity of BRCT mutants remained unchanged in the studies by Rodriguez et. al., suggesting that these mutants may participate in cytoplasmic functions independent of degradation ^29^. Santivasi et al. proposed that cytoplasmic localization of BRCA1 may facilitate lung metastasis in breast cancer ^62^, indicating that BRCA1’s cytoplasmic localization could contribute not only to the loss of nuclear DNA double-strand break repair functions but also to other oncogenic activities in the cytoplasm, such as centriole replication ^75^. We identified several target proteins with enhanced affinity for BRCA1-Y1853ter, validated through repeated identification across two distinct MS methodologies and further confirmed by structural modeling predictions of PPIs, supporting the robustness and reliability of these interactions. The correlation between *BRCA1* mutations and cancer incidence is evident in both homozygous and heterozygous states. These specific interactions may result from mutant BRCA1’s dominant negative effect on WT BRCA1 binding to its targets or from a stronger affinity that highlights unique interactions, thereby providing mutant BRCA1 with functions distinct from WT BRCA1 and promoting oncogenesis. These observations suggest a potential gain-of-function effect for mutant BRCA1. Further studies are needed to validate these interactions and determine their association with specific BRCA1 domains, with mechanisms likely involving the impact of BRCT domain mutations on the overall molecular conformation ^42^.

Numerous BRCT region sequence variations have been described, including nonsense and missense variants, of which 50% are VUS (variants of uncertain significance). The remainder are established mutations that have been observed to share various commonalities in aspects such as altered subcellular localization, PPI patterns, altered stability, and changes in drug resistance and transcription activation, as reported ^73,76–83^. Hence, these established mutants may collectively contribute to changes in BRCT function, potentially underlying shared pathogenic mechanisms, representing a valuable avenue for further investigation.

## Acknowledgments

We would like to thank the members of the Jensen lab for their critical review of this manuscript. Research reported in this publication was supported by the National Cancer Institute Cancer Center Support Grant P30CA168524 and the National Institute of General Medical Sciences of the National Institutes of Health R35GM145240 to M.P.W. We acknowledge the Integrative Imaging Core Laboratory, which is sponsored, in part, by NIH S10OD032207 at the University of Kansas Medical Center. The proteomics data were collected in the Mass Spectrometry and Proteomics Core facility utilizing the Orbitrap Ascend Tribrid System that was purchased with funds provided by the University of Kansas Cancer Center, which is supported by the National Cancer Institute Cancer Center Support Grant P30 CA168524.

## Author Contributions

Conceptualization: J.C., M.P.W., and R.A.J.; Molecular biology techniques and protein purification: J.C.; Mass spectrometry and analysis: M.J.R. and Z.S.C.; Data analysis and bioinformatics: J.C., M.P.W., M.E.S., J.N., and S.A.; Writing, original draft: J.C.; Writing, review and editing: M.E.S., M.J.R., M.P.W., and R.A.J.; Supervision: R.A.J. and M.P.W.; Funding acquisition: M.P.W. and R.A.J.

---

**Figure S1.**
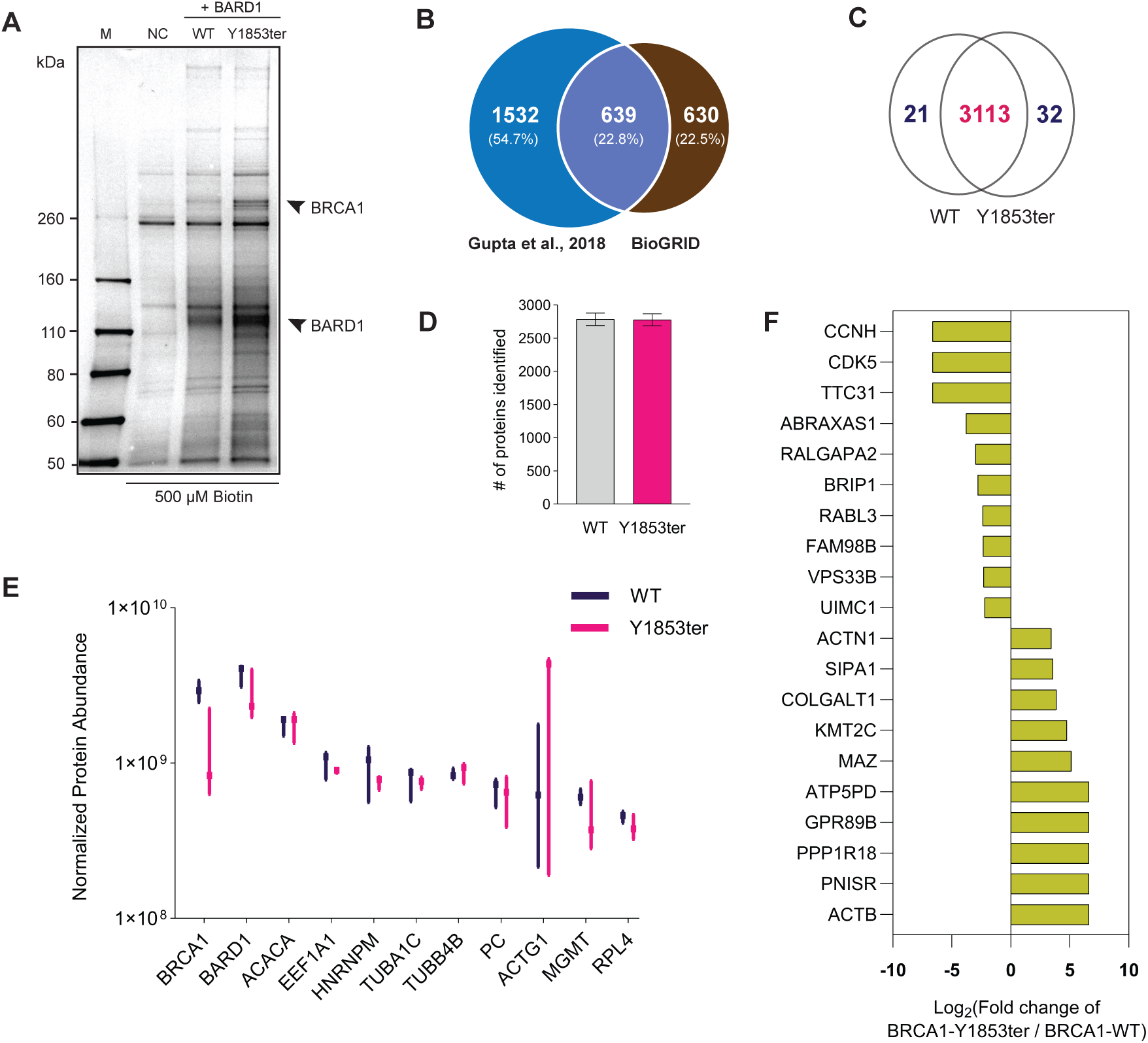
Cancer-specific interactome identified by TurboID-based MS. **(A)** Streptavidin-enriched eluates were analyzed by silver stain. 293T cells expressing TurboID constructs with either WT BRCA1 or BRCA1-Y1853ter, in the presence of 500 μM biotin labeling for 10 minutes, were subjected to streptavidin-based enrichment. Untransfected cells served as a control. Silver staining was used to visualize enriched proteins. **(B)** 54.7% of the identified PPIs in the studies of Gupta et al.^41^ are not represented in the BioGRID BRCA1 interactome database. **(C)** The Venn diagram illustrates the overlap of PPIs in WT and mutant samples. **(D)** Number of proteins identified in WT and mutant samples across three biological replicates. **(E)** Comparison of the top 10 highest abundance hits in WT and mutant samples across three biological replicates. **(F)** Ratio changes of the top 10 hits with the most significant upregulation or downregulation from the filtered list used for the volcano plot.

**Figure S2.**
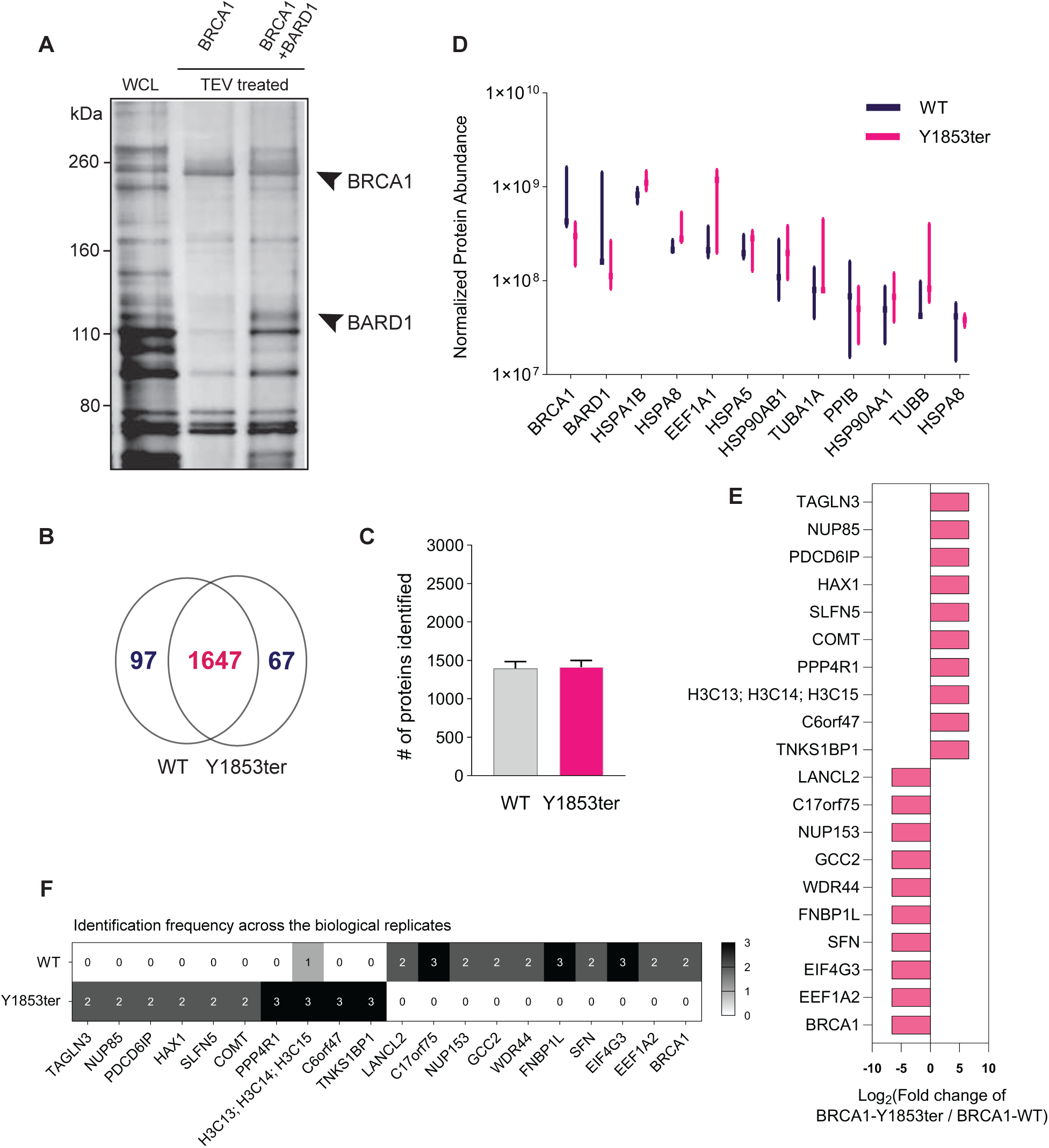
Cancer-specific interactome identified by Halo-based MS. **(A)** Purified eluates were subjected to silver staining analysis. 293T cells expressing either the WT BRCA1 Halo fusion construct alone or co-expressing BARD1 were analyzed. Whole cell lysate (WCL) prior to purification served as a control. Silver staining was employed to visualize and identify the enriched proteins. **(B)** The Venn diagram illustrates the overlap of PPIs in WT and mutant samples. **(C)** Number of proteins identified in WT and mutant samples across three biological replicates. **(D)** Comparison of the top 10 highest abundance hits in WT and mutant samples across three biological replicates. **(E)** Ratio changes of the top 10 hits with the most significant upregulation or downregulation from the filtered list used for the volcano plot. **(F)** The identification frequency of each listed individual across the three biological replicates of WT and mutant samples.

**Figure S3.**
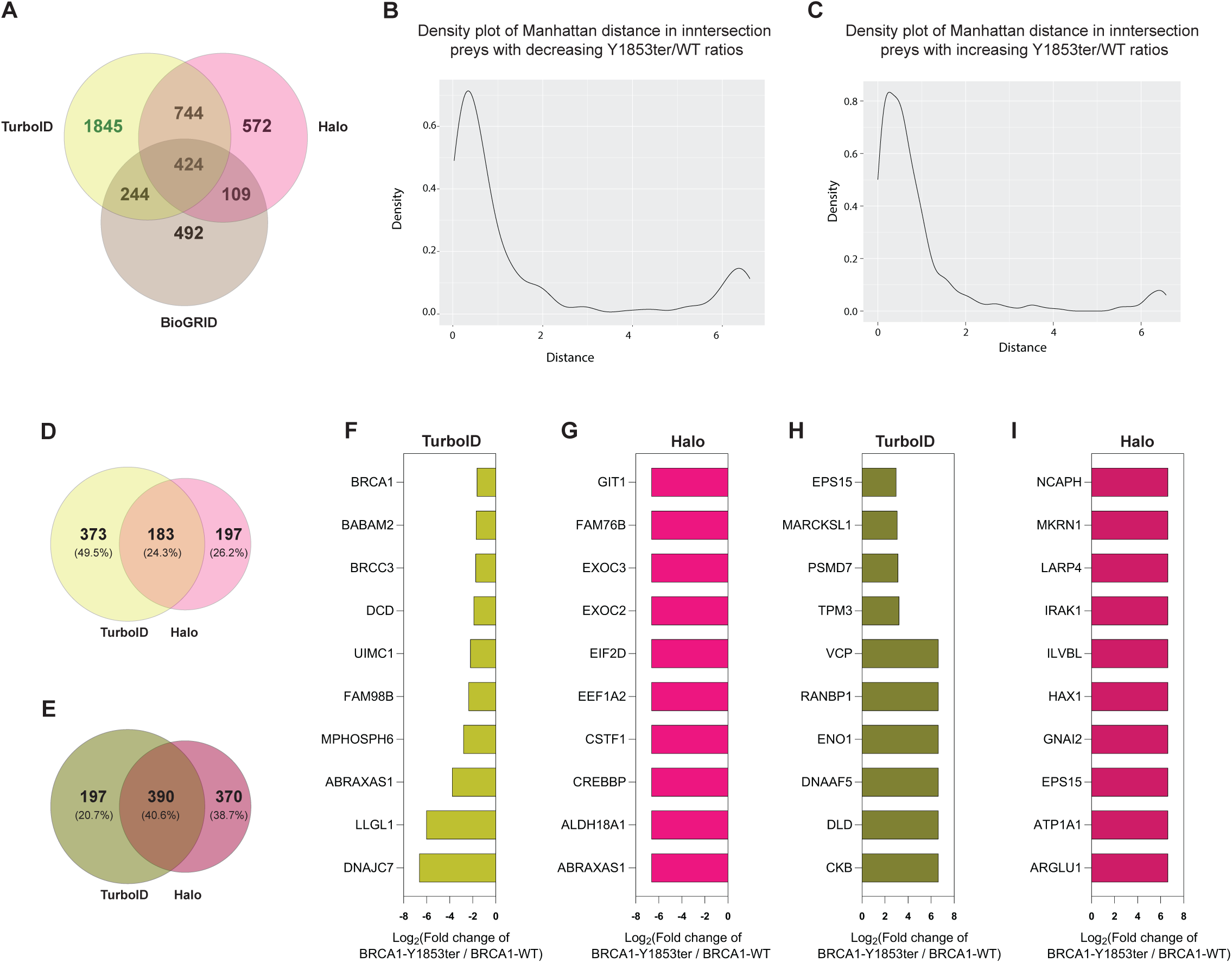
Venn diagram and statistical comparison of overlapping and method-specific hits. **(A**) Venn diagram illustrating the overlap of BRCA1 PPIs identified by the two methods employed in this study and those in the BioGRID BRCA1 interactome. **(B)** The density plot illustrates the distribution of Manhattan distance-based protein log_2_ fold changes between TurboID and Halo, specifically using a common subset of 183 upregulated proteins shown in (B) and 390 downregulated proteins shown in **(C)**. The plot depicts that most proteins have a minimal variance between the experiments (d<2), while a few proteins demonstrate a substantial difference (d>2). **(D)** Among the intersection preys, 183 hits demonstrate a consistent upregulation trend across both methods (D), while 390 hits exhibit a uniform downregulation trend **(E). (F)** Among the 183 hits, the top 10 hits with the most significant ratio changes identified by the TurboID method. **(G)** Among the 183 hits, the top 10 hits with the most significant ratio changes identified by the Halo method. **(H)** Among the 390 hits, the top 10 hits with the most significant ratio changes identified by the TurboID method. **(I)** Among the 390 hits, the top 10 hits with the most significant ratio changes identified by the Halo method.

**Figure S4.**
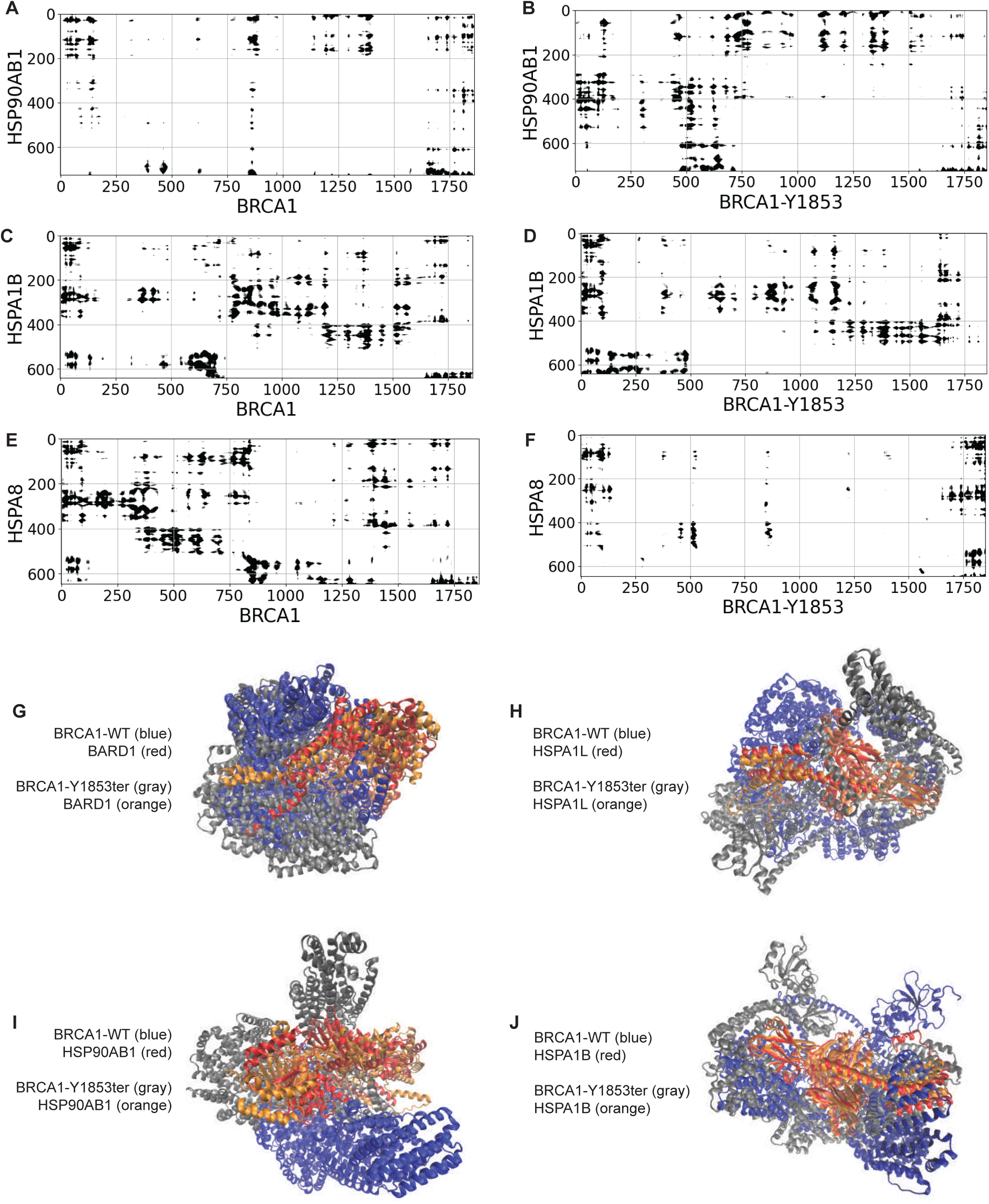
AF3-based comparative analysis of HSPs binding to WT and BRCA1-Y1853ter. **(A-F)** Contact map of WT and mutant BRCA1 interactions with HSPs. **(G-J)** Superimposed structural models illustrate alterations in the conformation and interaction patterns of BRCA1 WT and mutant upon binding to different preys.

**Figure S5.**
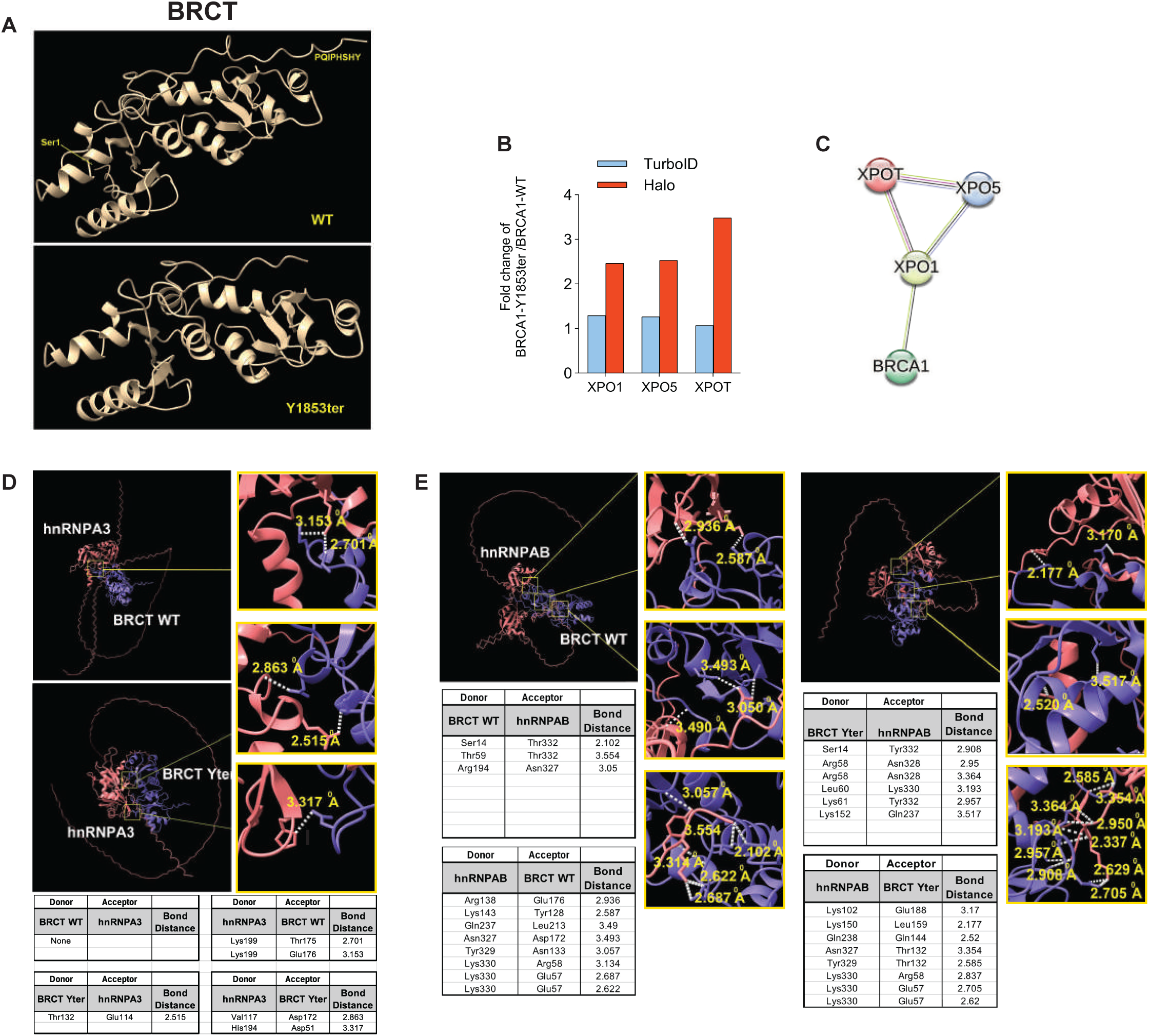
ChimeraX-based analysis of hnRNP interactions with BRCA1-Y1853ter and XPO family candidates. **(A)** Structural analysis of the BRCT domain in WT and mutant BRCA1. **(B)** Variation in the levels of several component proteins identified associated with nuclear export across two methods. **(C)** STRING highlights interactions and predicts connections between these transporter proteins and BRCA1. **(D-E)** ChimeraX-based structural analysis of WT and mutant BRCA1 interactions with hnRNPs.

**Figure S6.**
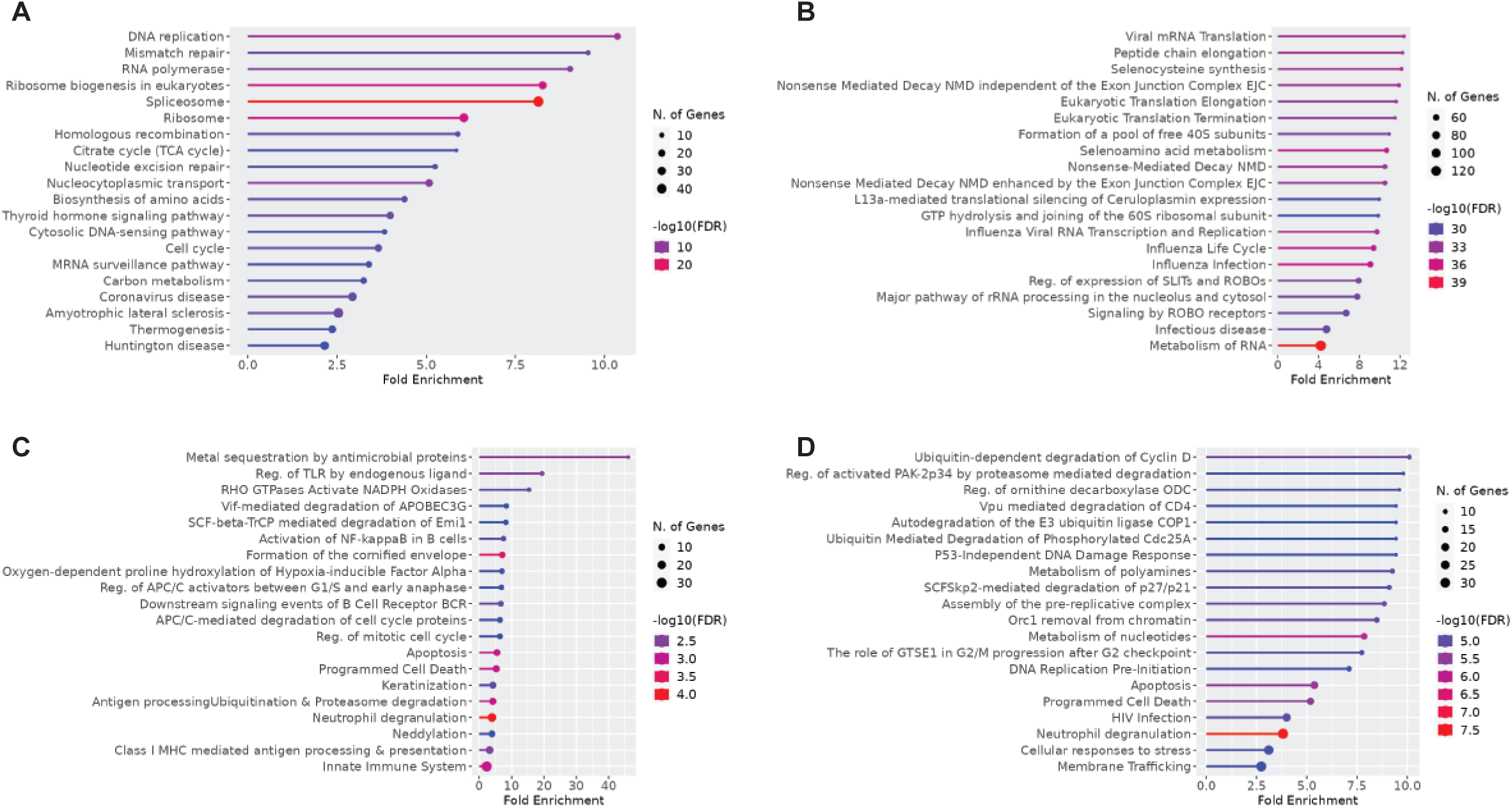
Unique PPIs from each method and the signaling pathway alterations predicted from these interactions. **(A)** TurboID-specific PPIs downregulated in the mutant compared to WT BRCA1. **(B)** TurboID-specific PPIs upregulated in the mutant compared to WT BRCA1. **(C)** Halo-specific PPIs downregulated in the mutant compared to WT BRCA1. **(D)** Halo-specific PPIs upregulated in the mutant compared to WT BRCA1. P-values reflect the statistical significance of enrichment, with lower values indicating a lower likelihood of the result occurring by chance (null hypothesis). False Discovery Rate (FDR) q-values adjust P-values for multiple testing to control type I errors. Fold Enrichment measures the magnitude of enrichment, with higher values indicating stronger enrichment and serving as an important metric of effect size. These results provide crucial insights into the impact of our research.

